# Enantiomeric histidine-rich peptide coacervates enhance antigen delivery to T cells

**DOI:** 10.1101/2024.09.10.612317

**Authors:** Ushasi Pramanik, Anirban Das, Elise M. Brown, Heather L. Struckman, Huihao Wang, Samuel Stealey, Macy L. Sprunger, Abdul Wasim, Jonathan Fascetti, Jagannath Mondal, Jonathan R. Silva, Silviya P. Zustiak, Meredith E. Jackrel, Jai S. Rudra

## Abstract

Peptides and peptidomimetics that self-assemble via LLPS have recently emerged as building blocks for fabricating functional biomaterials due to their unique physicochemical properties and dynamic nature. One of life’s most distinctive signatures is its selectivity for chiral molecules and, to date, coacervates comprised of D-amino acids have not been reported. Here, we demonstrate that histidine-rich repeats of (GHGXY)_4_ (X=L/V/P) and their enantiomers undergo LLPS opening new avenues for enhancing coacervate stability. Through a series of biophysical studies, we find that LLPS kinetics, droplet size, fusion, and encapsulation efficiency are dictated by the primary sequence. Further, these coacervates can encapsulate therapeutic cargo which are then internalized via endocytic mechanisms. Finally, we show that the coacervates enhance antigen presentation to CD4^+^ and CD8^+^ T cells resulting in robust proliferation and production of functional cytokines. Collectively, our study describes the development and characterization of enantiomeric peptide coacervates as attractive vaccine delivery vehicles with tunable physicochemical properties.

**HIGHLIGHTS:**

- D amino acid-peptides were used for the first time to construct phase separating coacervates
- Chirality does not restrict LLPS or modulate other coacervate properties
- Antigen delivery using chiral coacervates enhances and prolongs presentation to T cells

**PROGRESS AND POTENTIAL:** Peptides can undergo self-assembly via liquid-liquid phase separation (LLPS) to result in solute-rich coacervates that can serve as biomaterials. Using histidine-rich peptide repeats, this work demonstrates that peptides composed of entirely D-amino acids can form functional coaceravtes. The kinetics of LLPS and bulk properties of the droplets can be controlled through simple amino acid substitutions. The coacervates, while immunologically inert, exert an adjuvanting effect and enhance antigen presentation to T cells leading to proliferation and functional cytokine production. The materials showcased here possess high translational potential for combined delivery of immunomodulators and antigens for vaccine delivery against infectious diseases or cancer. The deliverables from this study will also inspire the development of chiral systems that will contribute to the knowledge of cellular processes associated with phase changes integral to both physiology and pathology.

## INTRODUCTION

Liquid-liquid phase separation (LLPS) is a phenomenon where associative interactions between macromolecules and segregative interactions with the external environment leads to the formation of solute-rich coacervates or droplets ^1–5^. LLPS promotes the spatiotemporal separation and concentration of vital cellular components, promoting processes including gene expression, proliferation, and differentiation ^6,7^. Intrinsically disordered proteins (IDPs) are particularly prone to undergoing LLPS. However, many natural and synthetic short peptides can also form simple or complex coacervates ^8–12^.

Peptides are attractive building blocks for biomolecular engineering due to the chemical diversity of amino acids. This diversity can also allow for modulation of key features that trigger phase separation and these features can also be modulated to tune the bulk properties of the resulting droplets. By tuning peptide primary sequences, the charge, hydrophobicity, and chirality of peptides can be modulated to achieve desired coacervate assembly-disassembly conditions, targeting to specific cells or tissues, and control over spatiotemporal release of loaded cargo ^13–16^. Compared to polymeric or inorganic nanocarriers that require organic solvents^17^, linkers^18^, and complex conjugation chemistry ^19^, droplets can form spontaneously in bioactivity-preserving aqueous buffers making them attractive for *in vivo* applications. Due to their dynamic nature, phase-separating droplets can undergo fusion with cell membranes leading to efficient transport of cargo into cells with minimal toxicity making them attractive for therapeutic delivery ^20^. These merits make peptide-based coacervates attractive functional biomaterials for cellular delivery of therapeutics ^21–24^.

A number of LLPS systems based on natural and synthetic peptides, as well as other block polymers have been described for applications as scaffolds for regenerative medicine and tissue engineering, bioactive drug delivery, and functional probes for disease diagnosis and therapy ^22–26^. In recent years, there has been an emerging interest in the introduction of chirality into biomaterials^27^. Chirality, inherent to all life processes, drives most biochemical reactions in organisms through selectivity for chiral molecular species (e.g., L-amino acids and D-sugars) ^28,29^. Owing to the natural use of L-amino acids, the fundamental change in backbone-side chain connectivity of D-enantiomers makes them resistant to proteases and immune recognition ^30,31^. To date, it is unknown how complete D-amino acid substitution affects kinetics of phase separation including droplet size, encapsulation efficiency, environmental sensitivity and stability; thus, can be a novel area of research.

Here we utilized LLPS-prone histidine-rich pentapeptide repeats of (GHGXY)_4_ (X=L/V/P) composed of all-L or all-D amino acids to assess how chirality affects coacervate formation. Our data shows that (GHGLY)_4_, (GHGVY)_4_, (GHGPY)_4_ and their enantiomers exhibit identical concentration-, pH-, and ionic strength dependent LLPS property. Substituting X with L/V/P resulted in different coacervate sizes of 1.8 µm/1 µm/0.2 µm, respectively. The encapsulation efficiency of various cargo (600 Da - 150 kDa) was also tested and rheological studies were conducted to assess viscoelastic behavior. Data showed that (GHGLY)_4_ coacervates were efficient at cytosolic delivery of DNA-encoding eGFP. Mechanistic studies showed that the droplets entered cells *via* energy-dependent endocytic pathways and microscopy data indicated that leucine droplets underwent fusion with other droplets as well as with cell membranes whereas valine droplets are resistant to fusion. On the contrary, valine coacervates augmented endo-lysosomal delivery of vaccine antigens for processing and presentation to T cells. Notably, coacervates of D-amino acids also led to increased and prolonged antigen presentation compared to their natural counterparts. Thus, our study demonstrates that LLPS are not modulated by chirality, and also exemplifies chiral phase separating systems are attractive vaccine delivery vehicles with tunable physicochemical properties.

## RESULTS AND DISCUSSION

### Phase separation of histidine-rich repeat peptides

The Miserez lab has pioneered the development of phase separating histidine-rich protein domains (HBPs) or peptides (HB*peps*) derived from the beaks of jumbo squid, demonstrating that they undergo LLPS to form coacervates, and have explored their potential application in biomedicine ^28^. Similar to VPGXG repeats in elastin, HB*peps* contain GHGXY repeats, where a minimum of four repeats are required for coacervate formation ^32^. Histidine and tyrosine residues are critical for phase separation of HB*peps*, which is driven by H-bonding of deprotonated histidine with tyrosine residues and proceeds in physiological buffers without the need for crowding agents^33^. The short length of HB*peps* and ease of production using synthetic chemistry facilitates the rational study of a range of constructs. To explore the role of chirality in HB*pep* coacervates formation, we introduced D-amino acids and other handles for post-synthetic modifications. Moduldation of chirality is a subtle change, in that enantiomeric peptides are identical in length, side-chain size, flexibility, hydropathy, charge, and polarity but allow for manipulation of certain biological outcomes. Here we generated all-L or all-D amino acid variants of the (GHGXY)_4_ peptide, where (X=L/V/P) and analyzed their coacervate properties along with utility as antigen delivery vehicles. To the best of our knowledge, this is the first report on the formation of coacervates composed of D-amino acids and the use of peptide coacervates for antigen delivery to T cells. The sequences of all peptides are shown in Figure 1A with lower case denoting D-amino acids. The identity and purity of the peptide variants was confirmed using MALDI-TOF (Figure S1, S2) and HPLC (Figure S3, S4).

**Figure 1.**
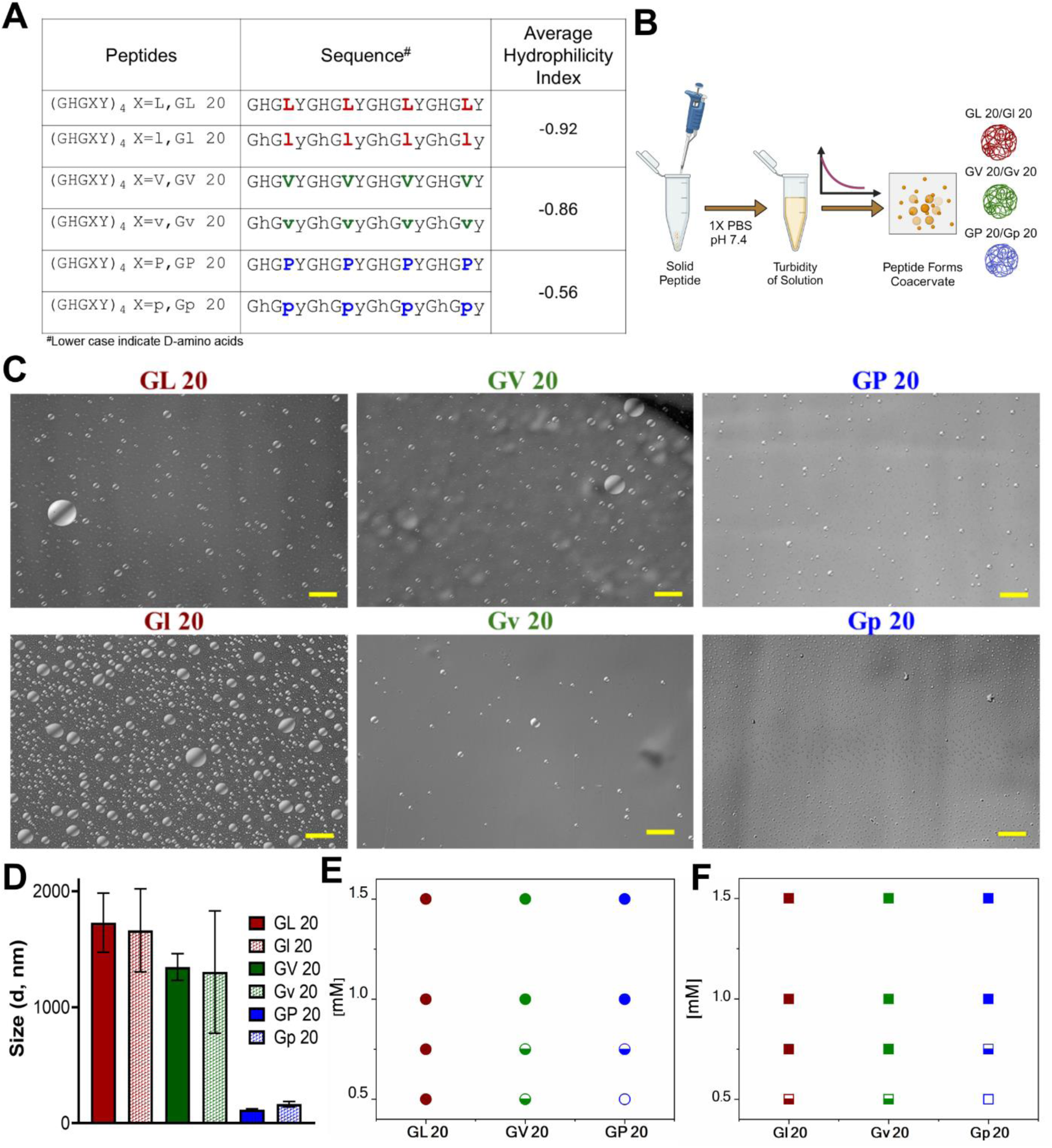
Chiral (GHGXY)_4_ sequences along with their LLPS properties. (A) Sequence of L- and D-peptides forming coacervates and their average hydrophilicity index (AHI). The AH index values were calculated using the online peptide calculator (https://www.bachem.com/knowledge-center/peptide-calculator/). Negative values indicate hydrophobicity. (B) Schematic of L- and D-peptide dissolution, followed by optical microscopy imaging. (C) Optical micrographs of chiral (GHGXY)_4_ peptides (scale: 50 µm). (D) DLS analysis shows size variation (diameter) among L- and D- (GHGLY)_4_, (GHGVY)_4_ and (GHGPY)_4_ (L>V>P). Data is an average (mean±SEM) of 3 technical replicates with 14 scans for each measurement. (E) and (F) Phase diagram of L- (e) and D- (GHGXY)_4_ peptides in a concentration-dependent manner. Filled circles/squares: complete coacervate formation; open circles/squares: no coacervate formation; half-filled circles: low coacervate formation.

### Chirality does not impact droplet characteristics or LLPS kinetics

The average hydrophilicity index (AHI) (https://www.bachem.com/knowledge-center/peptide-calculator/) was calculated for the variants and a negative value is indicative of their hydrophobic nature (L>V>P). Phase separation was tested using optical microscopy and we showed that no coacervates were formed in pure water or salt solutions (150 mM NaCl) (Figure S5, S6). The peptides formed coacervates only in physiological buffers (1×PBS, pH 7.4) (Figure 1B, C) and droplets were also visualized by TEM (Figure S7). DLS analysis showed that droplet sizes varied with hydrophilicity, but were identical for each pair of enantiomers and ranged from ∼1.8 mm for (GHGLY)_4_, ∼1.0 mm for (GHGVY)_4_, and ∼0.2 mm for (GHGPY)_4_ (Figure 1D). Prior studies using GHGXY analogs with multiple length repeats, variants, and substitutions have enhanced our understanding that hydrophobic interactions play a significant role in coacervation ^34^. Substituting a GAGFA repeat in HB*pep*s also led to the formation of dense hydrogels instead of liquid coacervates, presumably due to enhanced hydrophobicity and stronger intermolecular interactions ^33,34^. Based on these observations, the critical concentration (C_crit_) of coacervate formation was assessed. Data indicated that C_crit_ values decreased with increasing hydrophobicity of the amino acid at the X position (L=0.5 mM, V=0.75 mM, P=1 mM) (Figure 1E, 1F). The enantiomeric variants had identical C_crit_ values as expected (Figure S8, Table S1).

LLPS is promoted by increased solvent entropy resulting from peptide desolvation and increased peptide conformational freedom ^35^. The rapid movement of peptides inside the droplets leads to maturation and coalescence. The kinetics of LLPS was measured by observing the turbidity of the solution at 600 nm. We observed divergent kinetics of formation and maturation for the different variants. Solutions of (GHGLY)_4_ formed a turbid solution instantaneously as evidenced by the high A_600_ value at t=0 (Figure 2A), which decayed with time, likely reflecting the rapid maturation and coalescence of droplets ^12,36,37^. A striking difference in LLPS kinetics (turbidity measurement) was observed for (GHGVY)_4_, which steadily increased over 24 h (Figure 2B). Interestingly, no obvious turbidity changes were detected with time for the (GHGPY)_4_ peptide, suggesting that the initial formation of droplets observed upon its instant dissolution in 1×PBS may be the rate limiting step for the formation of coacervates (Figure 2C) ^37^. This difference in kinetics is likely due to hydrophobicity and inter- and intramolecular interactions that drive coacervate formation ^36,38,39^. We suppose that (GHGLY)_4_ has more propensity to form droplets instantaneously owing to its more hydrophobic nature, which also attributed to its rapid coalescence to reduce interfactial tension, resulting in the rapid decrease in turbidity measurements. Owing to (GHGVY)_4_ being less hydrophobic than (GHGLY)_4_, we observed a steady increase in the turbidity which can be a signature of slower droplet formation. As expected, all enantiomers exhibited kinetically identical profiles of LLPS (Figure 2A-C).

**Figure 2.**
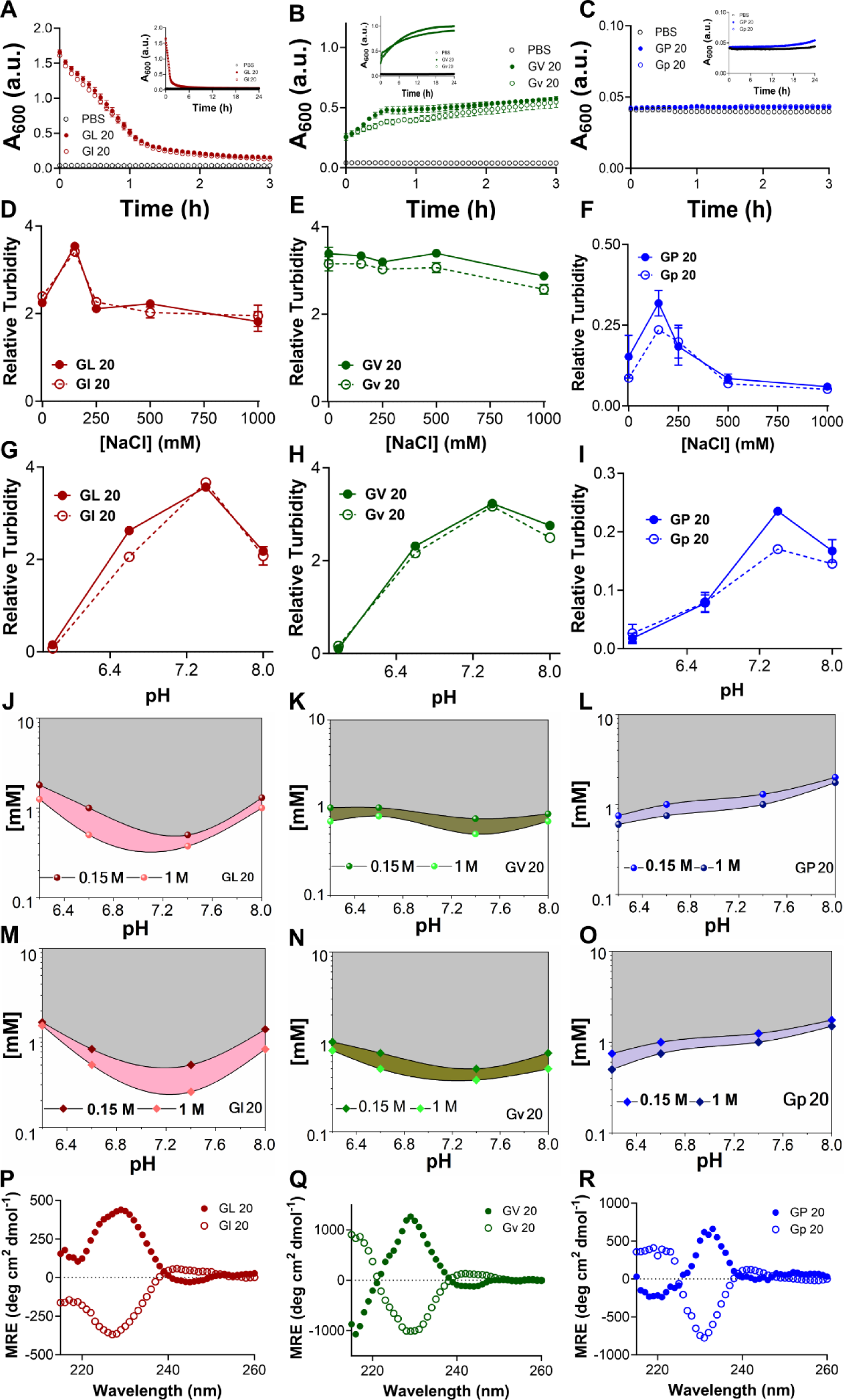
Phase separation of chiral (GHGXY)_4_ sequences with variation of pH and ionic strength along with their LLPS kinetics over a period of 24. **h.** Turbidity (λ = 600 nm) measured over time (3 h) for L- and D-peptides forming coacervates for (A) GL 20, (B) GV 20, (C) GP 20. Inset shows the full kinetics for 24 h for each peptide. Filled circles: L-peptides; open circles: D-peptides. Grey circles represent the control (only 1×PBS) to confirm specificity of the observed changes in peptide-induced turbidity. Relative turbidity (λ = 600 nm) of L- and D-peptides according to varied salt concentrations for (D) GL 20, (E) GV 20, and (F) GP 20. Turbidity measurements (λ = 600 nm) of L- and D-peptides at various pHs at 150 mM NaCl for (G) GL 20, (H) GV 20, and (I) GP 20. Critical phase separation concentrations of (GHGXY)_4_ variants *vs.* pH at 0.15 M and 1 M NaCl for (J) GL 20, (K) GV 20, (L) GP 20 (M) Gl 20, (N) Gv 20, and (O) Gp 20. CD profiles of 0.75 mM L- and D-peptides- (P) GL 20 (Q) GV 20, and (R) GP 20 in 1×PBS.

To further exclude the possibility that the drop in turbidity for the leucine peptides could be due to the settling of coacervates, turbidity measurements were made with or without agitation; resulting data indicated no differences between the two conditions (Figure S9). An acknowledged caveat with using turbidity assays to describe the kinetics of LLPS is that, plate readers are not fast enough to resolve early nucleation and growth events in droplet formation ^37^. As an additional measure of rigor and in accordance with LLPS reports, turbidity measurements were also carried out at 350 nm with no detectable differences (Figure S10). Overall, the stark differences in the kinetics of (GHGXY)_4_ variants demonstrates that amino acid hydrophobicity (L>V> P) drastically affects coacervate formation and maturation.

### The effects of temperature, salt concentration, and pH on coacervate formation

External factors, such as pH and ionic strength play an important role in phase separation by charge screening and ‘salting out’ effects. To test this, we assessed the effect of salt concentration on coacervate formation. All variants formed droplets within a wide range of NaCl concentrations (Figure 2D–F) but a narrow range of pH values (Figure 2G–I). The highest relative turbidity was observed at 150 mM NaCl and pH 7.4. This is not surprising as the peptides have a calculated isoelectric point of ∼7.66, where electrostatic repulsions are at a minimum and promote coacervation ^40^. Optical micrographs of coacervates under various salt and pH conditions are shown in Figures S11 and S12, respectively. Data confirmed the presence of coacervates in leucine and valine variants even in 1M NaCl whereas no coacervates were detected for the proline variant at >250 mM NaCl (Figure S11).

A complete understanding of the effects of salt on LLPS remains an outstanding challenge as phase separation depends on both the salt concentration and the chemical identity of ions called the Hofmeister effect ^41–43^. Wu *et al.* tested the effects of different ions of the Hofmeister series on HB*peps* and reported a broader two-phase region in the more kosmotropic sodium sulfate (Na_2_SO_4_) compared to sodium chloride (NaCl) or the more chaotropic sodium bromide (NaBr), which disrupted coacervation and allowed phase separation to occur only at higher peptide concentrations ^34^. It has been proposed that solubility at high salt concentrations is determined by the competition between the solvation energy and translational entropy of the ion ^41^. At higher salt concentrations, increasing hydrophobicity resulted in a significant decrease in C_crit_ (L>V>P) with a broader two-phase region for leucine peptides (Figure 2J, 2M). In contrast, enantiomers of valine (Figure 2K, 2N) and proline (Figure 2L, 2O) showed a comparatively narrow two-phase region. This is presumably due to shielding effect on protonated histidine residues, with reduced double-layer repulsion and entropy loss for counter-ion balancing, which in turn lowers C_crit_ for all variants ^44,45^.

To assess differences in temperature-dependent LLPS behavior of (GHGXY)_4_ peptides, turbidity measurements were made at room temperature (RT) and 37°C. The turbidity of the leucine peptide solutions decayed slightly faster at 37°C compared to RT, probably due to increased solubility. We detected very minute differences in turbidity for all peptides and their enantiomers at 37°C or RT (Figure S13). This is promising, as these properties could enable the encapsulation, retention, and delivery of therapeutics without significant concerns for their *in vivo* stability, which stands as a major goal for the use of these peptides.

### Secondary structures of peptide coacervates

We next probed the secondary structures of (GHGXY)_4_ variants using CD and FT-IR spectroscopy. The CD profiles of all peptides depicted ellipticity centered between 225 and 230 nm, attributed to π-π* transitions arising from aromatic interactions, also known as cotton effect (Figure 2P-R) ^46,47^. Moreover, we observed a red-shift in ellipticity maximum from 228 nm (GHGLY)_4_ to 230 nm (GHGVY)_4_ to 232 nm (GHGPY)_4_, consistent with a decrease in hydrophobicity from L>V>P, known as positive cotton effect ^47^. The CD spectra of all enantiomers were mirror images of each other as expected. FT-IR spectra of the variants showed the subtle differences in their secondary structures (Figure S14). The amide-I bands of leucine and valine peptides were centered around 1643 and 1647 cm^−1^, respectively. The valine peptide had an additional band at 1634 cm^−1^. In contrast, proline variant had bands centered at 1628 and 1637 cm^−1^, which typically represent *β*-sheet structures. However, none of the peptides exhibited *β*-sheet fibril formation and all-atom molecular simulations found minimal to no *β*-sheet content in all variants (Figure 4M).

### Mechanical properties of peptide coacervates

It is now widely recognized that biomolecular coacervates can be viscoelastic and studies on IDP domains have reported both shear-thinning or shear-thickening behavior ^48,49^. We assessed the mechanical properties of freshly prepared coacervate solutions using parallel plate rheology at a fixed shear rate (Figure 3A). Data indicated that their viscosity was dependent on peptide hydrophobicity (L>V>P), where we observed leucine droplets exhibited greater viscosity compared to valine or proline variants (Figure 3B). Increasing the peptide concentration from 1 to 2 mM led to an increase in viscosity for all peptides (Figure 3B). Prior reports on LLPS by various peptides have reported an environment-dependent shear thinning or shear thickening behavior and increasing the shear rate led to a decrease in viscosity with no differences observed between the enantiomers (Figure 3C). Such shear-thinning behavior may be advantageous for the use of coacervates as injectable biomaterials. Further, we measured the storage and loss moduli (*G*′ and *G*″, respectively, Figure S15) of the coacervate solutions and the ratio of G′/G″>1 indicated viscoelastic nature. Leucine coacervates exhibited nearly a 3-fold increase in *G*′ compared to valine droplets, whereas a 2-fold increase in *G*′ values were noted for valine compared to proline. Rheological studies on minimal HB*pep* sequences have highlighted that the viscoelastic properties of the coacervates could be tuned by single residue mutations which can in turn alter the secondary structure content ^34^. The higher storage modulus of coacervates has been attributed to increased *β*-sheet content ^34^. While no secondary structures have been observed for the (GHGXY)_4_ variants, their mechanical properties strongly correlated with hydrophobicity and our data support prior findings that the viscoelastic properties of coacervates can be tuned by small changes in the peptides’ primary sequences.

**Figure 3.**
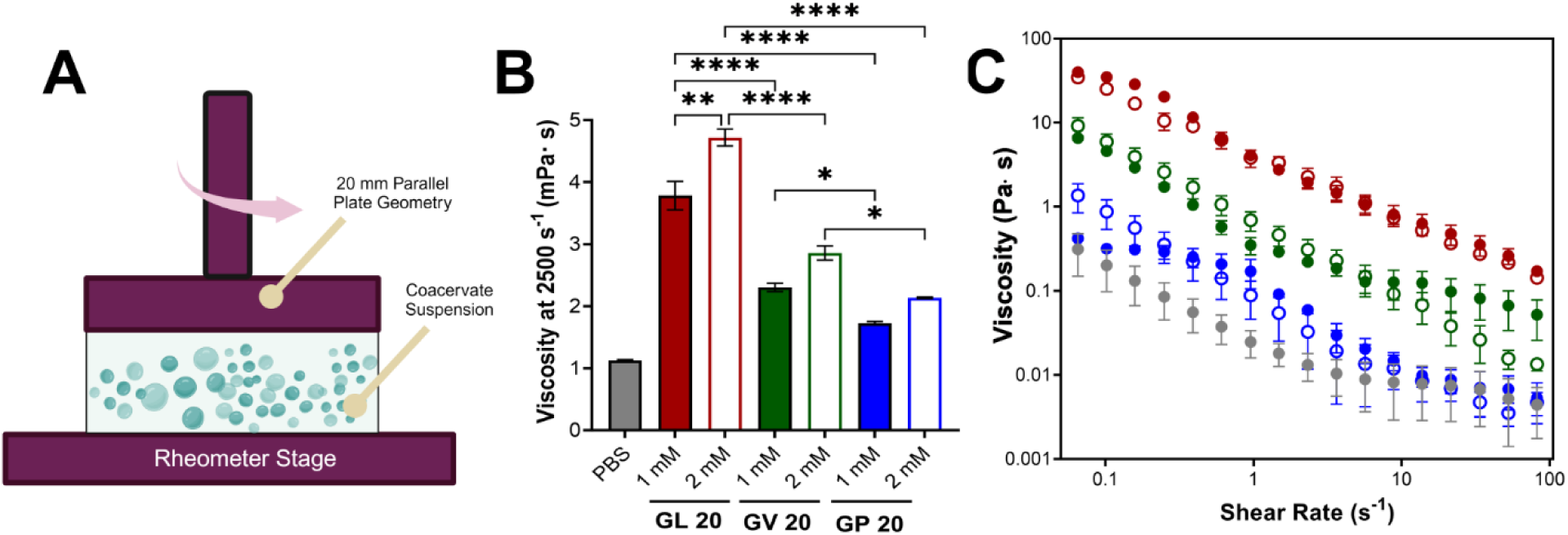
Mechanical testing of peptide coacervates. (A) Schematic of coacervate testing using a rotational rheometer. (B) Viscosity measured at a shear rate of 2500 s^−1^ using a Hagen-Poiseuille viscometer. Data is representative of three biological replicates and plotted as mean±SEM. (C) Measured viscosity at varying shear rates using a rotational rheometer at a peptide coacervate concentration of 1 mM. The filled and open circles represent L- and D-GL 20 (red), GV 20 (green), GP 20 (blue), respectively. Grey circles represent 1×PBS alone. *p < 0.05, **p < 0.01, ****p < 0.0001 as determined by a one-way ANOVA.

### Deciphering molecular interactions by all-atom simulations

To understand the molecular interactions driving coacervate formation, we conducted multi-chain, all-atom simulations of (GHGXY)_4_ variants, where we started the simulations from a dispersed state (Figures 4A-C) and ended up with a single droplet for each variant during the simulations (Figures 4D-F). We next examined the average inter-peptide contact probabilities for each peptide (Figure 4G-I). Each contact probability was calculated as a time average of the residue-residue contacts between any two chains, subsequently averaged over the number of peptide pairs. This contact map was generated after all the chains in the simulation box have merged into a single droplet, providing an overview of the molecular interactions within each chain that drive and stabilize coacervate formation. We observed that the pattern in the contact map varied between different peptides, despite 80% sequence homology. This demonstrates that even a single residue change in the repeating unit of the sequence can result in very different interactions driving LLPS, potentially contributing to distinct rheological and physical properties of the phase-separated droplets. Notably, all sequences showed high contact probabilities between histidine residues, although differences in histidine-tyrosine interactions varied widely between peptide sequences. This suggests that (GHGXY)_4_ peptides tend to form coacervates via strong and frequent cation-π and π-π interactions.

**Figure 4.**
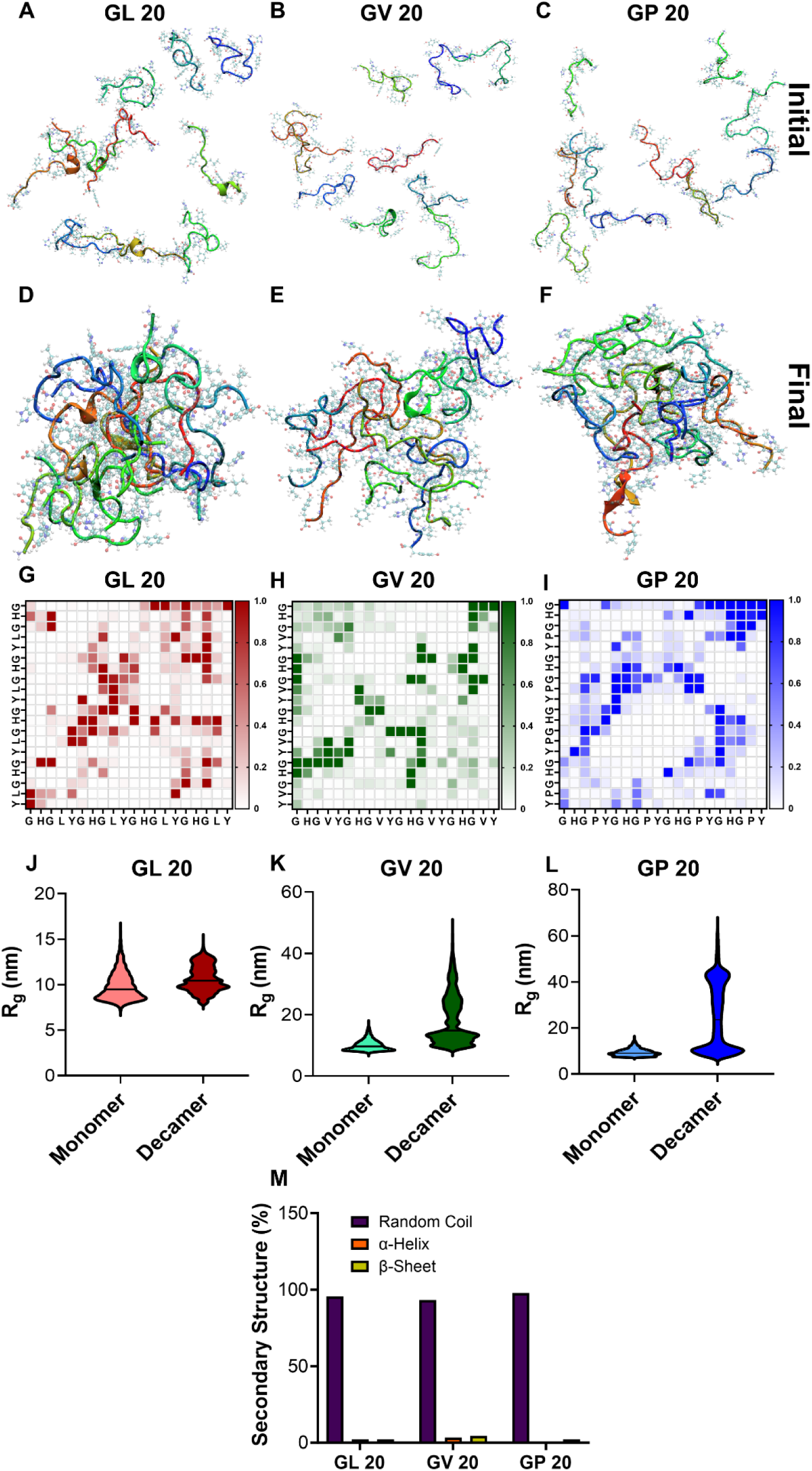
All atom simulations depict molecular interactions in LLPS by (GHGXY)_4_ peptides. Initial and final snapshots of multichain all-atom simulations for (A, D) GL 20, (B, E) GV 20, (C, F) GP 20 considering 10 monomeric peptides for each inside a box. Contact map for (G) GL 20, (H) GV 20, (I) GP 20 generated upon merging of all the chains in the simulation box into a single droplet. This contact map provides an overview of the molecular interactions within each chain responsible for driving and stabilizing LLPS by these peptides. The colour scale represents lowest (0, white) to highest (1, colour) contact probability between the peptide amino acids in 2 of such peptides (horizontal and vertical axes). R_g_ plots confirming LLPS for the monomer and decamer of (J) GL 20, (K) GV 20, (L) GP 20 (M) The percentage secondary structure calculated from all-atom simulation for (GHGXY)_4_ peptides.

To quantify these interactions, we calculated the number of π-π interactions during the last 500 ns of the simulation trajectory, a period before which all chains have formed a single, stable droplet. We calculated the radius of gyration (R_g_) of individual chains under two scenarios: (i) a single chain in a box of water and (ii) a chain in a droplet. We then plotted the corresponding R_g_ values for both cases (Figures 4J-L) and observed an increase in R_g_ for scenario (ii), a signature of LLPS ^50,51^. We next characterized the π-π interactions responsible for driving droplet formation; data in Figure S16 A-F provides relative snapshots of parallel and perpendicular π-stacking interactions from the simulation trajectories. We also compared the number of parallel and perpendicular π-stacking interactions that occurred during the last 500 ns of simulation trajectories for all three peptides. (GHGPY)_4_ showed the highest number of parallel π-stacking interactions, followed by (GHGLY)_4_ and (GHGVY)_4_ (Figure S16G). A similar trend was observed for perpendicular π-stacking interactions (P>L>V). These findings support data in panels Figure 4G-I, where (GHGPY)_4_ showed high contact probabilities between H and Y, compared to L or V variants.

### Encapsulation efficiency and cellular uptake of coacervates

We next tested the capacity of (GHGXY)_4_ coacervates to encapsulate cargo for cellular delivery. The encapsulation efficiency was calculated from absorbance or fluorescence measurements of the cargo in the dilute and the condensed phases and confirmed using microscopy (Figure 5A, S17). Using eGFP, we calculated the loading efficiency to be the highest for the leucine droplets (∼90%) compared to valine (∼75%) or proline (∼30%) coacervates (Figure 5B). A wide range of cargo (600 Da-150 kDa) were tested and data confirmed that encapsulation efficiency varied with hydrophobicity (L>V>P) (Figure S18). Our findings differ from reports by the Lampel lab wherein the most hydrophobic peptide droplets had the lowest GFP encapsulation efficiency while the most polar peptide droplets entrapped the greatest quantity of GFP ^38^. A plausible explanation can be that the number of π–π, cation–π, or hydrogen bonding interactions between the peptide and cargo could dictate the loading capacity, thereby explaining the differences in encapsulation between the peptides ^38^. Due to their small size and poor encapsulation efficiency, (GHGPY)_4_ droplets were not tested further.

**Figure 5.**
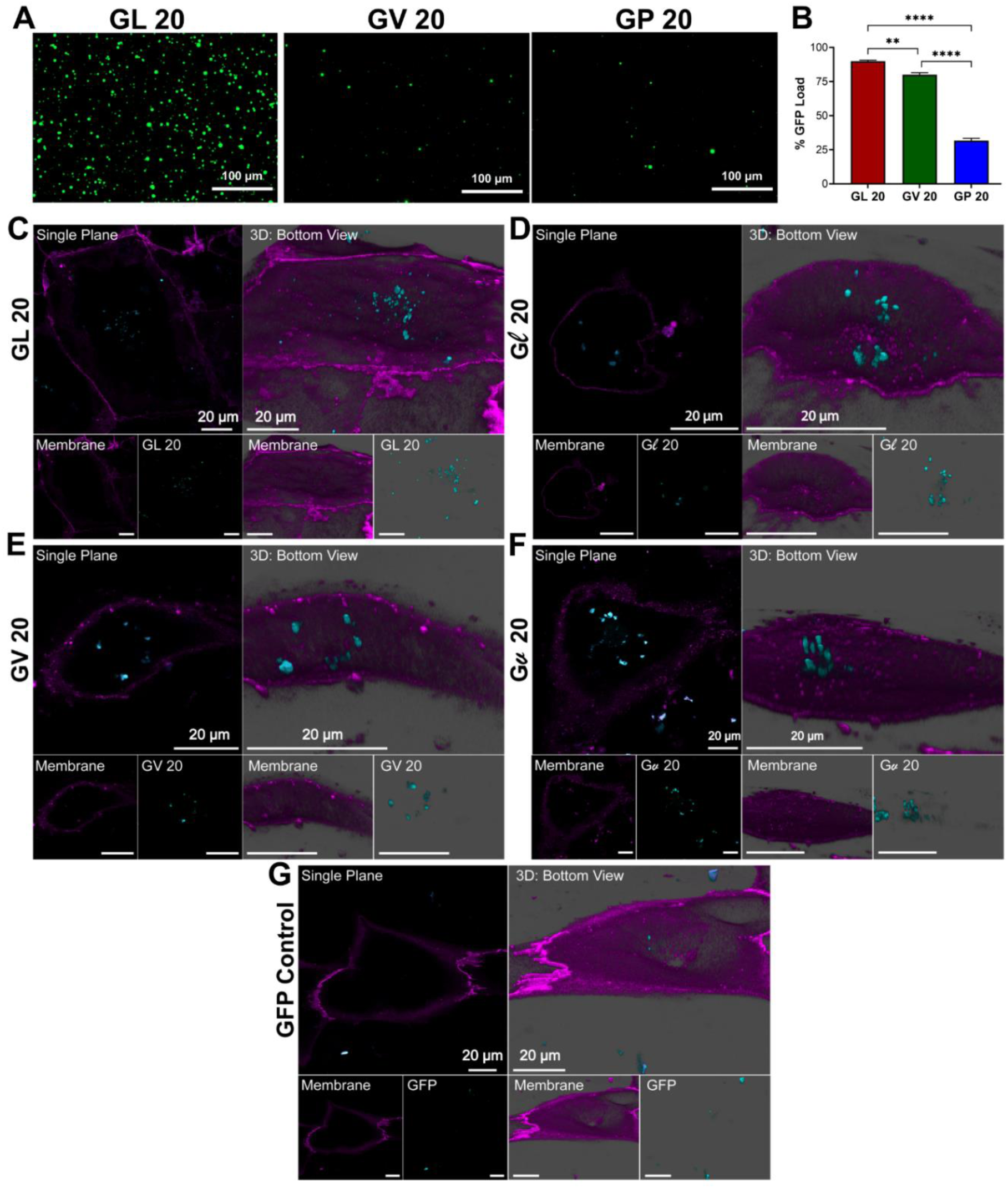
Localization of GL 20, Gl 20, GV 20, Gv 20 within hiPSC-CMs. Fluorescence microscopy images of eGFP loaded (A) GL 20, GV 20, and GP 20 coacervates (Scale bar: 100 µm). (B) % eGFP load by GL 20, GV 20, and GP 20 droplets. **p < 0.01, ****p < 0.0001 as determined by a one-way ANOVA. Representative confocal images from hiPSC-CMs treated with (C) GL 20, (D) Gl 20, (E) GV 20, (F) Gv 20, (G) eGFP control (teal). The cellular membrane was identified with WGA membrane stain (purple). Micrographs were presented as single plane and 3D snapshots from the bottom view perspective (Scale bars: 20 µm).

In order to further illustrate the the utility of these peptide forming coacervates as delivery vehicles in a wide variety of cell lines, we next used eGFP loaded (GHGLY)_4_ and (GHGVY)_4_ coacervates and confocal microscopy to visualize cellular uptake in hiPSC-CM (human induced pluripotent stem cell-derived cardiomyocytes) cultures. After 30 min of incubation, single plane bottom views confirmed their localization inside the cells (Figure 5C–F). Images show that both leucine and valine coacervates (teal) are fully encapsulated by the membrane (purple) of the hiPSC-CMs and are centrally located. Qualitatively the coacervates improved the delivery efficiency compared to the soluble eGFP control (Figure 5G). Single plane top view images were also acquired to confirm intracellular localization of the droplets (Figure S19). No significant differences were noted between the internalization of the enantiomeric variants. These results demonstrate that the encapsulation efficiency of (GHGXY)_4_ coacervates can be tuned by the chemical composition of the peptide building blocks; further, leucine and valine variants can efficiently deliver the entrapped cargo to cells *in vitro*.

### The coalescence properties of coacervates

Similar to water, fusion of biomolecular coacervates into larger droplets has been reported for various membraneless organelles including nucleoli, stress granules, and Cajal bodies ^52^. Dynamic coacervates can undergo rapid fusion despite the high macroscopic viscosity inside them. Given the differences in hydrophobicity, size, viscoelasticity, and encapsulation efficiency between the coacervates, we next tested their fluidity by capturing video still images over time (Figure 6A, B). Data indicated multiple rapid fusion events for the leucine coacervates only which agrees with the LLPS kinetics data in Figure 2A, where a rapid decay in turbidity was attributed to droplet fusion (video in Figure S20). However, the droplet edges did not fully resolve suggesting arrested fusion mechanisms ^53,54^. As an additional readout, individual leucine coacervates loaded with red, blue, or green fluorophore tagged proteins were mixed (red: PE-Cy7-mouse IL-4; blue: EF 450-mouse IFN-γ; green: FITC mouse anti-mouse H-2K^b^) and coalescence was observed over time (Figure 6C). Microscopy data showed droplets with secondary colors (fusion of two primary colors) or grey (fusion of three primary colors) suggesting droplet fusion (Figure 6D), after about 15 minutes of mixing. This was also validated by increased leucine droplet size over time (Figure S20). In comparison, valine droplets did not exhibit fusion behavior (Figure 6G, H). To test whether the coacervates can encapsulate multiple cargo molecules without exclusion or compositional drift, we mixed RBG fluorophore tagged proteins prior to loading (Figure 6E, I). Data showed that the bulk of the leucine and valine droplets encapsulated all three proteins with few coacervates showing only binary mixtures (Figure 6F, J). Control experiments with mixed protein solutions without coacervates were used to validate encapsulation within the droplets (Figure S21).

**Figure 6.**
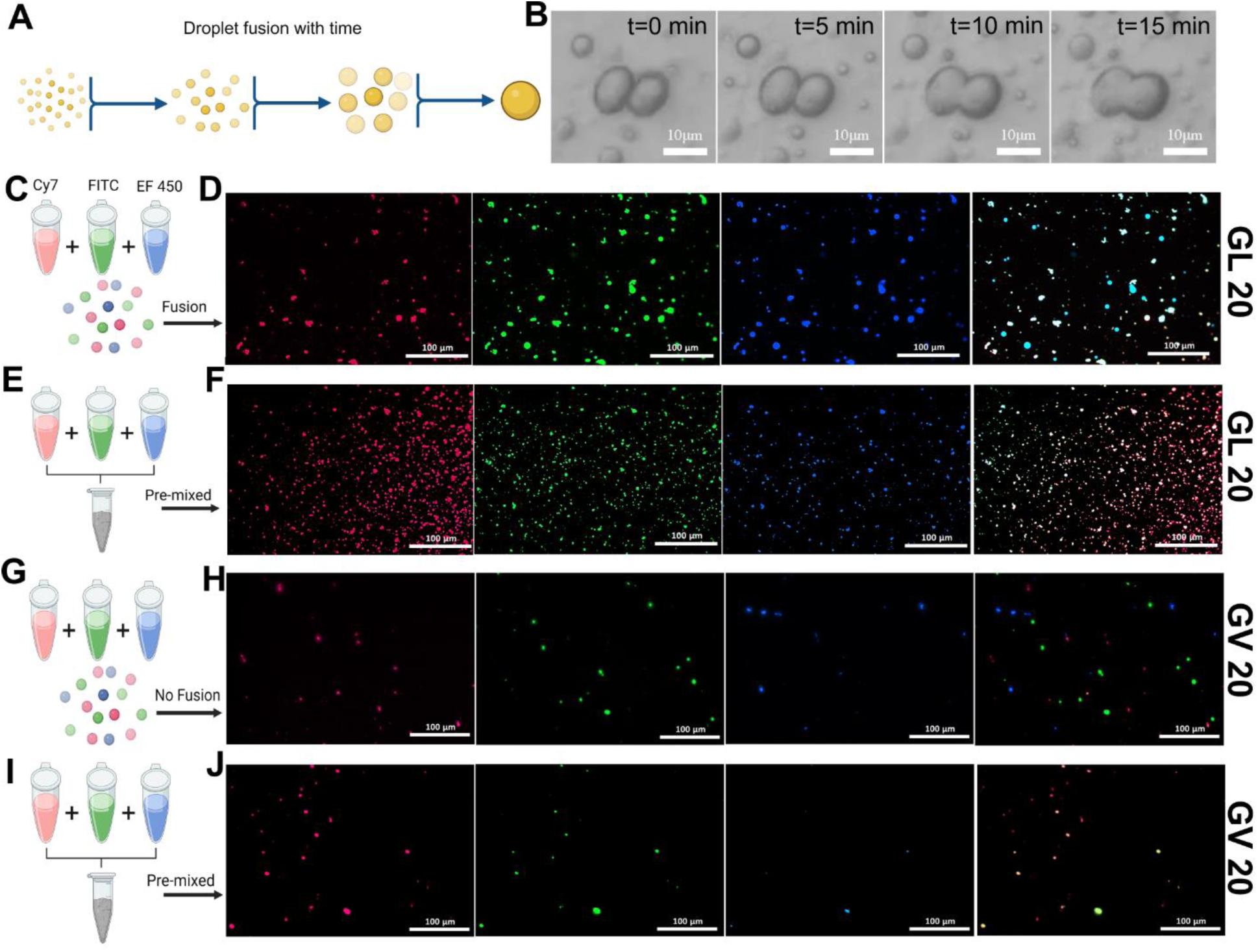
Droplet coalescence over time. (A) Schematic of droplet coalescence over time to generate larger droplets. (B) Sequence of enhanced snapshots extracted from a continuous video recording that captured the dynamic coalescence events of peptides GL 20 over 15 min. Each snapshot, selected from frames captured every 500 milliseconds, highlights pivotal moments in the coacervation and coalescence process. These images show the coalescence event of the phase-separating peptides as they transition from isolated droplets to interconnected coacervates. The snapshots are timestamped to document the progression and timing of each significant event, providing a detailed visual narrative of peptide behavior under experimental conditions. Schematic of fluorescently labeled cargo loaded in (C) GL 20 (G) GV 20 coacervates and subsequent coalescence over time. (D) Red, blue, green, and overlayed channels depicting GL 20 coalescence over time, producing grey droplets. Addition of individual labeled protein solutions in (E) GL 20 peptide (I) GV 20 to observe cargo uptake by GL 20 and GV 20 peptide coacervates. (F) Red, blue, green, and overlayed channels show GL 20 droplets do not preferentially load different cargos and all three proteins can be loaded in one (grey) droplet. (H) Red, blue, green, and overlayed channels show GV 20 peptides do not coalesce over time and are present as single-colored droplets. (J) Red, blue, green, and overlayed channels show GV 20 droplets do not preferentially load different cargos. Most GV 20 peptides could load two proteins in one droplet, owing to their smaller size.

At the interface of the coacervate-bulk phase, the surface molecules experience a net force toward the interior of the bulk phase. This interfacial tension drives fusion of two encountering liquid-like coacervates. A recent study by Sun *et al.* reported that fluid-like or gel-like coacervates can be achieved *via* systematic hydrophobic or charged amino acid mutations in GHGXY repeats ^55^. The study also found differences in uptake rates and intracellular release kinetics of cargo where fluid-like droplets exhibited enhanced cell membrane adhesion and wetting compared to gel-like coacervates ^55^. Our data suggests that the leucine coacervates have a low to moderate interfacial tension and undergo fusion readily compared to the valine droplets, thereby decreasing its interfacial area through droplet coalescence ^39^. Using confocal microscopy, we also observed that leucine coacervates interacted with and crossed the cell membrane (orange dashed circle, Figure 7A, 7B left panel). The same droplet was found to be deformed and stretched spanning the intercellular, membrane, and extracellular spaces as shown in the orthogonal projection (right side Z plane panel) (Figure 7C). Also, intracellular leucine coacervates were found to contain membrane signal (yellow dashed circle) indicating coacervate-lipid interactions. Colocalization of membrane signal (purple) with coacervates (teal) in the intracellular space of hiPSC-CMs is shown in Figure 7B. These observations suggest that the leucine droplets traverse the cell membrane, and these events may play a role in their internalization. Despite our best efforts, this behavior was not observed for valine coacervates.

**Figure 7.**
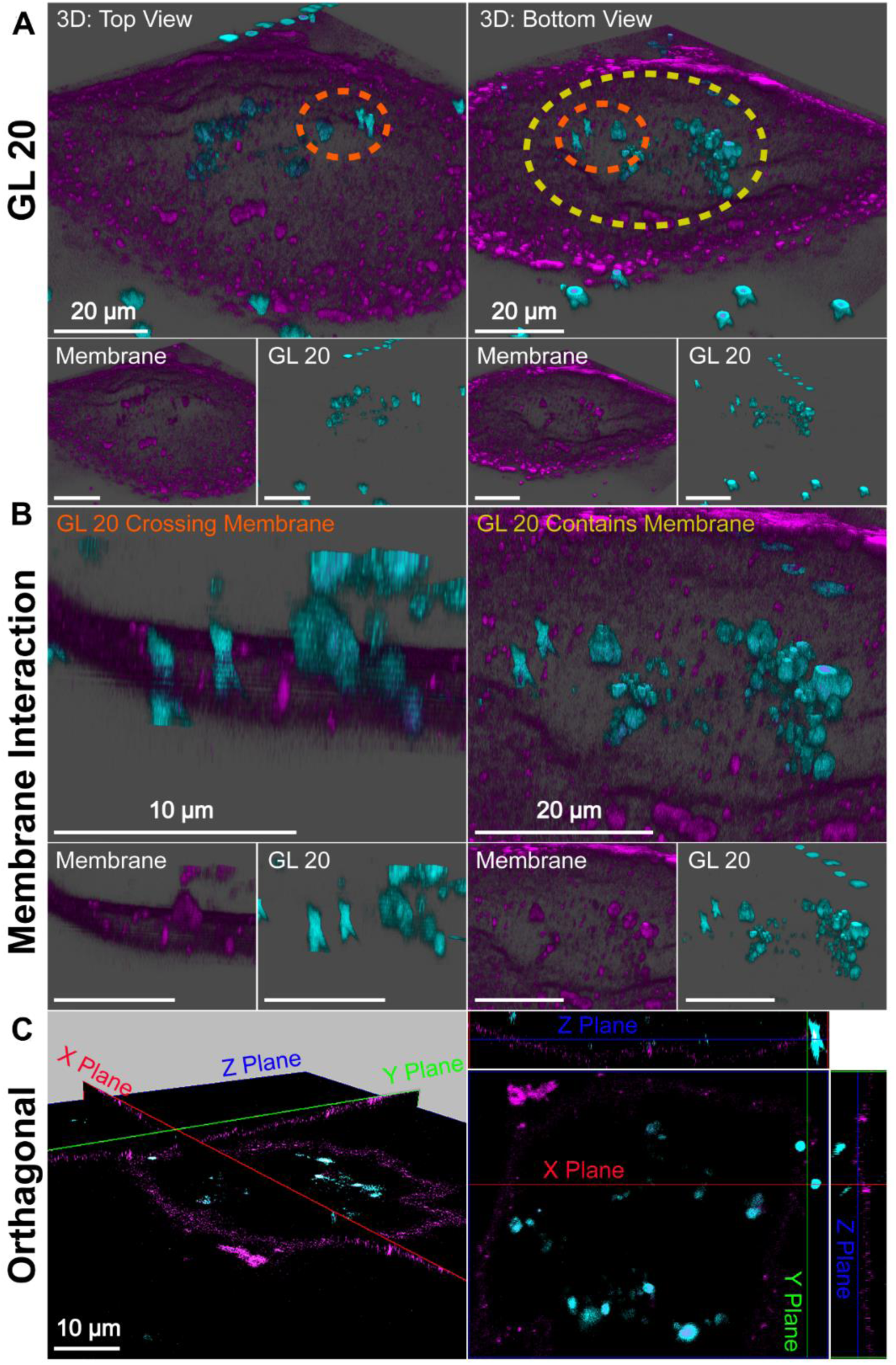
Membrane interactions of GL 20 in hiPSC-CMs. Representative confocal micrographs of hiPSC-CMs treated with GL 20 (teal). The cellular membrane was identified with WGA membrane stain (purple). (A) Micrographs are presented as 3D snapshots from the top and bottom views (scale: 20 µm) (B) Membrane interactions of GL 20 were captured as transmembrane crossings (orange, scale: 10 µm) and colocalization with the membrane (yellow, scale: 20 µm) (C) Orthogonal projections in the X-, Y-, and Z-planes (scale: 10 µm) further captured GL 20 transmembrane crossing.

We next tested whether the fluid-like properties of leucine coacervates enable efficient cytoplasmic delivery of cargo using droplets loaded with eGFP plasmid DNA. Data indicated robust eGFP expression in HEK cells treated with the leucine droplets (Figure S22). Qualitatively, the delivery efficiency was lower compared to Lipofectamine 2000 but higher compared to the naked plasmid or valine droplets. Poor eGFP expression was detected in cells treated with valine coacervates and we hypothesize that due to their lower interfacial tension, these droplets are presumably sequestered in endosomes. The coacervates were not cytotoxic under the conditions tested (Figure S23, S24).

### The mechanisms of cellular internalization of coacervates

To test mechanisms of internalization, we utilized *in vitro* antigen presentation assays (Figure 8A). Here, bone marrow derived dendritic cells (BMDCs) are treated with coacervates, which are loaded with the model antigen ovalbumin (OVA). Following incubation, BMDCs are washed extensively to remove extracellular coacervates and overlaid with hybridomas that specifically recognize the OVA_323-339_ peptide in the context of MHC class II (DOBW cells) or the OVA_257-264_ peptide in the context of MHC class I (CD8 OVA 1.3 cells). Secretion of IL-2 in response to peptide-MHC recognition is an indirect measure of coacervate uptake and can be quantified using ELISA (Enzyme-Linked Immunosorbent Assay) (Figure 8A). In DCs treated with OVA loaded (GHGLY)_4_ and (GHGVY)_4_ coacervates at 4°C and overlaid with DOBW cells, significantly lower IL-2 levels were detected compared to 37°C controls (Figure 8B, D). To further confirm that the differences in uptake were not due to reduced plasma membrane fluidity, we used ATP-depleted media at 37°C to block all energy-dependent pathways (Figure 8C) ^56^. Again, a significant loss of IL-2 secretion was observed suggesting that the droplets are internalized through energy-dependent mechanisms (Figure 8D). A key advantage of this assay compared to flow cytometry is that it eliminates positive readouts from membrane-bound coacervates and requires complete internalization and processing of the cargo. We next used pharmacological inhibitors of endocytic mechanisms to explore mechanisms of cell entry (Figure 8E). BMDCs were treated with inhibitors of clathrin-mediated endocytosis (Chlorpromazine, CPZ)^57^, dynamin dependant endocytosis (Dynasore, Dyn) ^58^, macropinocytosis (Wortmannin, Wort) ^59^, cholesterol (Methyl β-CD) ^60^, and IPA3 the regulatory domain of PAK1 (IPA-3) ^61^ for 1 h prior to antigen presentation assays. Treatment with Wort had the most significant effect on coacervate uptake, signifying that formation of macropinosomes likely plays important role in their endocytosis. In contrast, CPZ, which disrupts the formation of clathrin mediated uptake, the GTPase inhibitor, Dyn, which inhibits dynamin activity leading to inhibition of endocytosis, Methyl β-CD, which disrupts cholesterol-mediated lipid rafting and inhibition of PAK1 signaling by IPA3, did not significantly impact droplet uptake (Figure 8F). This is similar to that observed by Miserez and group, who observed energy dependant micropinocytosis mechanism as the process by which the HB*pep* and HB*pep*-SP peptides are delivered inside the model membranes ^62^.

**Figure 8.**
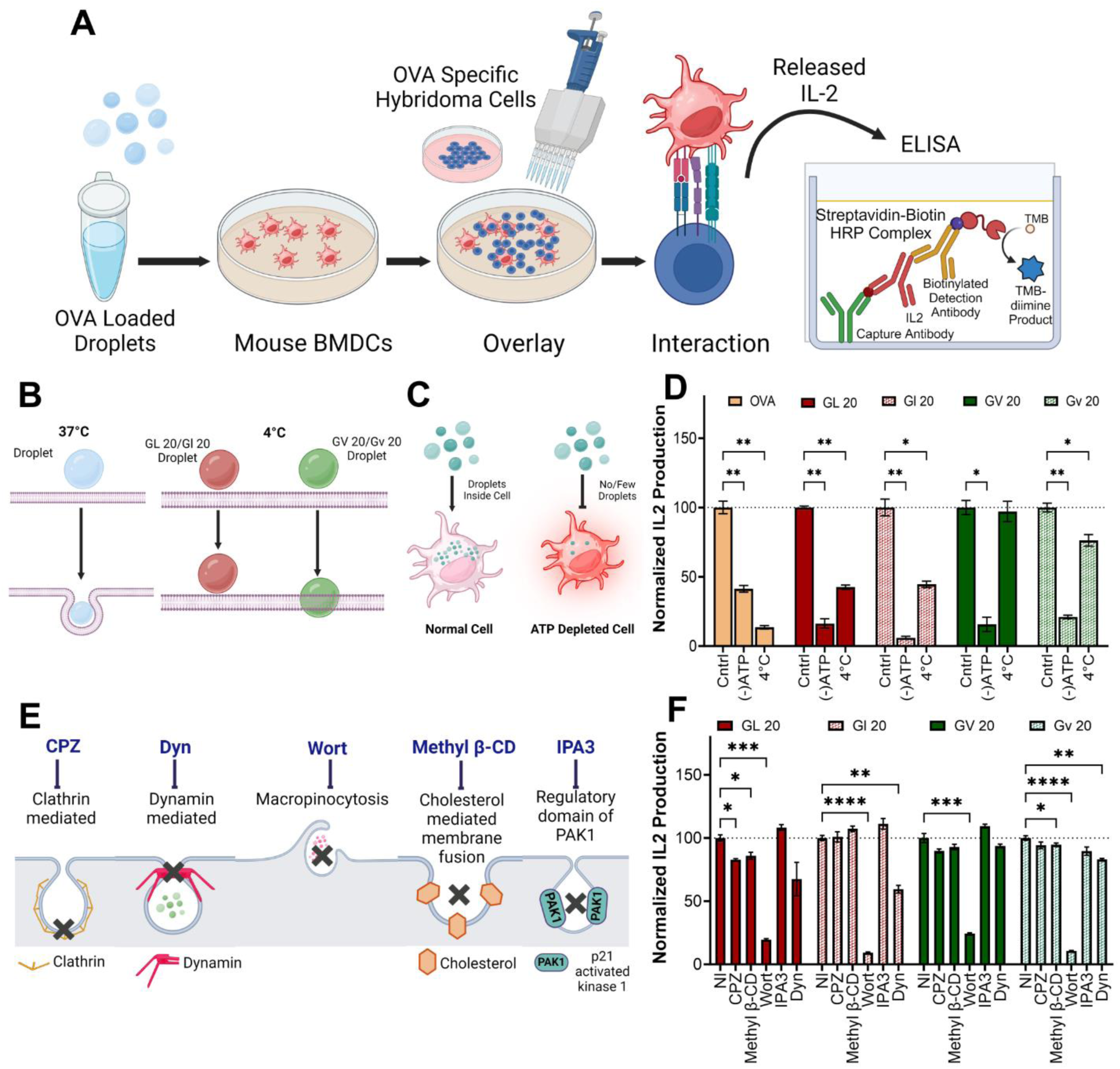
Mechanistic study entry of (GHGXY)_4_ droplets in mouse BMDCs. (A) Schematic of ELISA to analyze IL-2 production induced delivery and processing of OVA in mouse BMDCs upon interaction with OVA specific Hybridoma cells mediated by chiral (GHGXY)_4_ coacervates. (B, C) Schematic representation of delivery of OVA loaded L- and D-peptides in ATP depleted cells, and at 4°C, depcted with respect to 37°C (Cntrl). (D) IL2 production by OVA loaded L- and D-peptides in presence of ATP depletion, and at 4°C, normalized with respect to 37°C (Cntrl). (E) Schematic of different endocytosis inhibitors and their target pathways. (F) Effect of endocytosis inhibitors on the uptake of L- and D-(GHGXY)_4_ (L/V) coacervates loaded with OVA in mouse BMDCS. A pronounced inhibitory effect was observed for Wortmannin (Wort), which inhibit macropinocytosis. \**p* < 0.05, \*\**p* < 0.01, \*\*\**p* < 0.001, \*\*\*\**p* < 0.0001 as determined by one-way ANOVA.

### Activation of innate immunity by coacervates

For the continued development of peptide coacervates as vaccine delivery vehicles, understanding their ability to activate innate immune cells is paramount. BMDCs are key regulators of adaptive immunity with the potential to induce T cell activation/immunity or T cell suppression/tolerance. Previous studies have demonstrated that peptide biomaterials can exert adjuvant-like effects by inducing the upregulation of maturation markers such as MHC II, CD80, and CD86 and the secretion of chemokines (MCP-1α/CCL2, KC/CXCL1) and cytokines (GM-CSF, IL-5, IL-6, IL-1β)^63,64^. To test this, we treated BMDCs with (GHGLY)_4_ and (GHGVY)_4_ coacervates or LPS as a positive control and assessed cytokine and chemokine production using multiplex assays. Data indicated that the coacervates were immunologically inert and did not induce the production of pro-inflammatory (IL-6 and TNF-α), tolerance-inducing (IL-10) or maturation-specific (IL-15) cytokines (Figure S25). We simultaneously profiled the expression of inflammatory chemokines namely: monocyte chemoattractant protein-1 (MCP-1), macrophage inflammatory proteins (MIP-1α/MIP-1β) and IFN-γ-inducible protein 10 (IP-10) which are crucial to mounting an effective immune response due to their ability to recruit other immune cells (Figure S25). No chemokines or growth factors (GM-CSF, EGF, MIF, and PGDF-BB) that modulate immune cell proliferation or host immune responses were detected, suggesting that the coacervates do not activate innate immune signaling. These purely structural scaffolds can now be advantageously modified with select immune agonists that activate pathways specific to conferring protection against infections, non-infectious diseases, autoimmune disorders, or allergies.

### Antigen delivery, processing, and presentation mechanisms

Dendritic cells scan peripheral tissues for antigens, which once acquired are carried to the lymph nodes where they induce adaptive immune responses. This requires the processing and breakdown of the antigens in lysosomes into peptide fragments that can be presented to T cells. A number of polymeric carriers exploiting lysosomal delivery and lysosomal escape have been reported for the induction of effective immune responses ^65^. To assess the ability of the coacervates to deliver antigens to lysosomal compartments, we loaded (GHGLY)_4_ and (GHGVY)_4_ with DQ-OVA which upon protease-mediated hydrolysis in the lysosomes produces fluorescent peptides. Green fluorescent puncta were detected in DCs treated with both coacervates, indicating lysosomal processing of DQ-OVA (Figure 9A, B, S26). To quantify delivery, we pre-treated DCs with the coacervates for 2 h, 4 h, 24 h, 48 h, and 72 h prior to hybridoma overlay and measured IL-2 levels in the supernatant (Figure 9C). Compared to soluble OVA, the coacervates enhanced antigen presentation and IL-2 production increased with time up to 24 h, decreased at 48 h, and was not detectable at 72 h. Importantly, for the same antigen dose, IL-2 production was significantly higher in DCs treated with (GHGVY)_4_ droplets compared to (GHGLY)_4_ (Figure 9C). Further, comparison of the enantiomers indicated prolonged antigen presentation by D-form (GHGVY)_4_ coacervates with robust IL-2 signal detectable even at 72 h compared to the L-form (Figure 9D). Surprisingly, this effect was absent in D-leucine treated cells. Taken together, these findings indicate that chiral peptide coacervates are excellent biomaterials for enhancing antigen delivery and prolonged presentation for applications in vaccine development. To confirm our findings with the model antigen OVA, we used Ag85B, an immunodominant protein from *Mycobacterium tuberculosis* and BB7 hybrodima cells, which recognize Ag85B_240-254_ in the context of MHC class II. Data indicated enhanced antigen presentation of Ag85B when delivered using (GHGLY)_4_ or (GHGVY)_4_ coacervates or their enantiomers (Figure S29).

**Figure 9.**
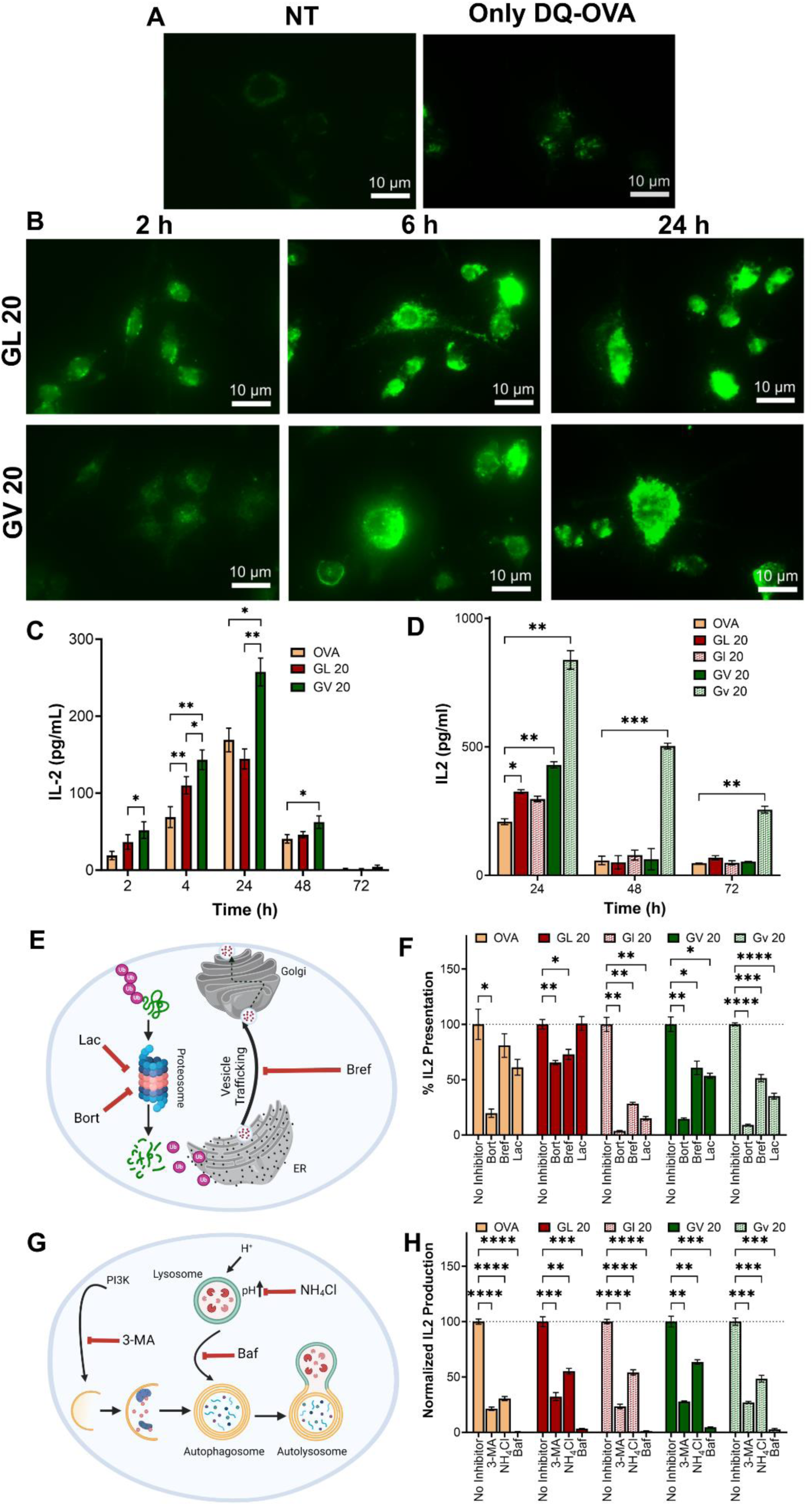
Delivery of the model antigen OVA and its fluorescent form (DQ-OVA) in mouse BMDCs mediated by (GHGXY)_4_ coacervates. (A, B) Time dependent DQ-OVA persistence and presentation inside mouse BMDCs with the aid of coacervates formed by (GHGXY)_4_ (L/V) as compared to no treatment and DQ-OVA only controls. (C, D) IL-2 production upon successful delivery and presentation of OVA-loaded (C, 1 µM; D, 10 µM) GL 20, GV 20 coacervates inside mouse BMDCs from 2 to 72 h vs. free OVA. Gv 20 shows persistent presentation of OVA over 72 h of treatment. (E) Schematic depicting inhibition of MHC-I antigen presentation by MHC-I specific inhibitors. (F) Inhibitory effects on IL-2 production by MHC I inhibitors (Bortezomib, Brefeldin A, and Lactacystin) after delivery of chiral (GHGXY)_4_ (L/V) loaded with OVA to mouse BMDCs. (G) Schematic depicting inhibition of MHC-II antigen presentation by MHC II specific inhibitors. (H) Inhibitory effects on IL-2 production by MHC II inhibitors (3-MA, NH_4_Cl, and Bafilomycin) after delivery of chiral (GHGXY)_4_ (L/V) loaded with OVA to mouse BMDCs. The cells were overlayed with Ova 1.3 cells (MHC I group) and DOBW hybridomas (MHC II group). \**p* < 0.05, \*\**p* < 0.01, \*\*\**p* < 0.001, \*\*\*\**p* < 0.0001 as determined by one-way ANOVA.

Classically, exogenous antigens internalized by DCs are processed in the lysosomal compartments and presented *via* MHC-II to CD4^+^T cells, whereas cytosolic antigens processed *via* the proteasomal pathway and presented *via* MHC-I to CD8^+^T cells. Non-canonical presentation of exogenous antigens on MHC-I and cytosolic antigens on MHC-II has been reported to confer protection against some viral infections and cancers. We next tested whether antigens delivered using peptide coacervates are processed *via* classical MHC-I (Figure 9E) or MHC-II pathways (Figure 9G). DCs were treated with the three different inhibitors of each pathway 30 min prior to coacervate addition, and IL-2 secretion by DOBW or OVA1.3 hybridoma cells were used to assess MHC class II or MHC class I antigen presentation, respectively. At the concentrations tested, none of the inhibitors were significantly cytotoxic (Figure S27). A significant decrease in IL-2 production was detected in DCs pre-treated with Bafilomycin, NH_4_Cl, or 3-methyl adenine (3-MA) which are well-known MHC II inhibitors (Figure 9H). Bafilomycin specifically inhibits V-ATPase, which prevents the acidification of endosomes and lysosomes and NH_4_Cl acts by raising the pH of intracellular compartments, thus disrupting the processing of endocytosed materials. Lastly, 3-MA is an autophagy inhibitor that blocks class III PI3K activity. Further, fluorescence intensity in DCs treated with DQ-OVA coacervates was diminished in the presence of NH_4_Cl or Bafilomycin, suggesting reduced proteolytic activity (Figure S28). Similarly, MHC class I presentation was significantly reduced as evidenced by lower IL-2 secretion in all groups treated with Bortezomib, Lactacystin, or Brefeldin A. Bortezomib and Lactcystin are proteasomal inhibitors, whereas Brefeldin A inhibits the trafficking of degraded peptides from the endoplasmic reticulum for MHC class I presentation (Figure 9F). Together, these findings suggest that antigens delivered using peptide coacervates are processed and presented *via* classical MHC-pathways.

### Induction of functional T cell responses by coacervates

While hybridoma cell lines serve as tools to quantify antigen presentation and elucidate mechanisms, T cell proliferation assays assess the ability of antigens to induce T cells to divide and produce effector cytokines as measures of activation. We used CD4^+^T and CD8^+^T cells isolated from transgenic mice that specifically recognize OVA_323-339_ and OVA_257-264_ epitopes in the context of MHC class II or MHC class I molecules, respectively. Purified T cells were stained with a fluorescent dye and overlaid onto DCs treated with OVA-loaded coacervates. Proliferation was quantified based on the fluorescence intensity of the daughter cells and cytokine production was assessed using multiplex assays (Figure 10A). Our data showed robust proliferation of both CD4^+^T and CD8^+^T with cells with no significant differences between groups or enantiomers for CD4^+^T cells (Figure 10B). However, the proliferation in case of (GHGLY)_4_ group was slightly higher than the (GHGVY)_4_ group. Similarly robust proliferation was detected in CD8^+^T cells and the percentage of proliferated cells was higher in the (GHGVY)_4_ group compared to the (GHGLY)_4_ group (Figure 10I). A significant difference was also noted between (GHGLY)_4_ enantiomers. To determine the nature of the T cell response, we measured the cytokine profile in the supernatants. Data showed robust levels of Th1 (IFN-γ and IL-2) and Th17 (IL-17) cytokine production by OT-II and OT-I T cells compared to Th2 (IL-4, IL-5) cytokines (Figure 10C-F, H-K, S30-S33). Th1 cytokines, particularly IFN-γ, are proinflammatory and are necessary for the control of intracellular viral infections, while Th17 cytokines (e.g., IL-17A) mediate host defensive mechanisms, especially against extracellular bacterial infections. These results pave the way for future work to establish efficacy of coacervates as effective vaccine delivery vehicles for combating bacterial or viral infections.

**Figure 10.**
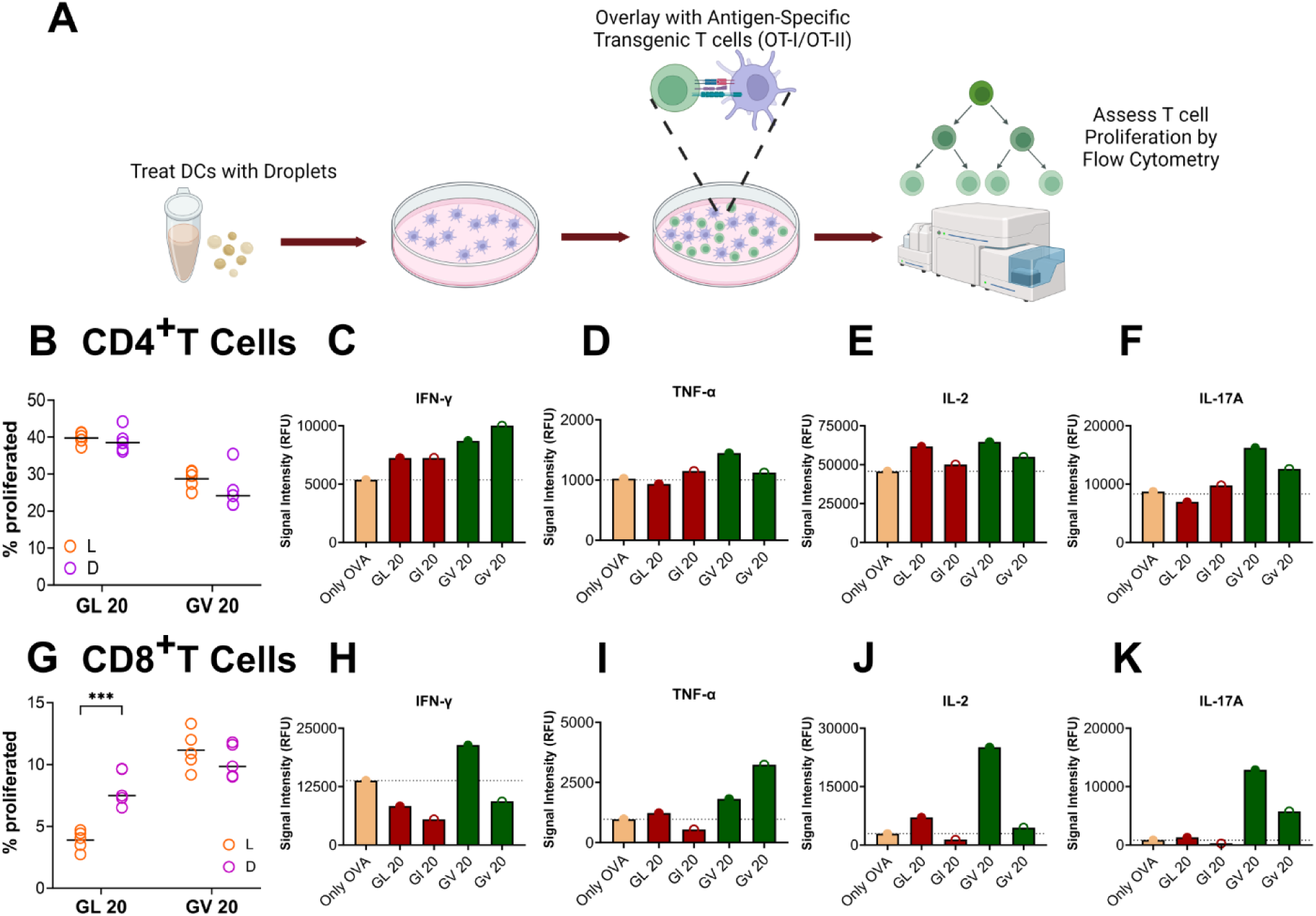
OT-I/OT-II mouse CD8^+^/CD4^+^T cell proliferation and cytokine release upon L- and D-(GHGXY)_4_ (L/V) mediated delivery of OVA. (A) Schematic representation of antigen specific transgenic T cells (OT-I/OT-II mice) on antigen presenting mouse BMDCs to assess T cell proliferation. (B) OT-II mice CD4^+^T and (G) OT-I mouse CD8^+^T cell proliferation upon treatment with L- or D- (GHGXY)_4_ coacervates loaded with OVA compared to control (additional data in Figure S30 and S32) Data are shown as mean ± SEM of 5 technical replicates. Effect of L- and D- (GHGXY)_4_ peptide coacervate-mediated OVA delivery on production of IFN-γ, TNF-α, IL-2, and IL-17A. Mouse BMDCs were treated with free OVA or coacervate-loaded OVA (10 µM) for 24 h before overlaying with CTV-labeled OT-II/OT-I hybridoma cells for 66 h followed by measurement of (C, H) IFN-γ, (D, I) TNF-α, (E, J) IL-2, (F, K) IL-17A in the culture supernatant of OT-II and OT-I mice, respectively. Values are shown as means of 2 wells in a Mouse Adaptive CodePlex Secretome chip pooled from treatments of 4 wells. \*\*\**p*< 0.001, as determined by 2-way Anova analysis.

## CONCLUSIONS

In conclusion, we have demonstrated the development and testing of enantiomeric peptide coacervates as biomaterials for antigen delivery and presentation to T cells. The D-amino acid coacervates reported in this study are a leap forward not only for the development of novel materials but also as model systems to further our understanding of the role of the peptide/protein coacervates in disease and health. The inter- and intramolecular interactions between peptides, the interfacial tension and fusion, and the bulk properties of the resulting droplets can be tuned via simple amino acid substitutions in the primary sequence. Studies on cargo encapsulation and cellular uptake mechanisms have verified that chirality is not a prerequisite for their use as delivery vehicles in biomedical applications. The ability to improve delivery to cytosolic or endosomal compartments based on the primary sequence enables one to control desired biological functions. The inert nature of the coacervates, combined with their ability to enhance antigen presentation, provides opportunity for their further modification with immunogenic or tolerogenic signals to control the immune response. Because of the spontaneous formation of the coacervates in a bioactivity-preserving physiological environment and the efficiency of antigen delivery, they hold great potential for the development of vaccines and immunotherapies against chronic infectious and non-infectious diseases.

## MATERIALS AND METHODS

### Peptide synthesis and purification

Peptides (GHGLY)_4_, (GHGVY)_4_, and (GHGPY)_4_ were synthesized using standard Fmoc-SPPS chemistry on Rink Amide with Oxyma (Ethyl cyano(hydroxyimino) acetate) and Diispropyl carbodiimide (DIC) as coupling agents on a Liberty Blue microwave-assisted synthesizer. Peptides were cleaved using a cocktail of trifluoroacetic acid (TFA), tri-isopropyl silane (TIS), and H_2_O (95:2.5:2.5) and extracted and washed in cold diethyl ether. The resulting pellet was dissolved in acetonitrile/water mixture (50:50) and frozen prior to lyophilization. Peptides were purified (∼90%) using High Performance Liquid Chromatography (HPLC) on a Dionex Ultimate 3000 HPLC equipped with a diode-array detector on an Agilent Poroshell RP-C18 column (4.6 mm×150 mm), operating at a flow rate of 1 mL/min. Peaks corresponding to 220 nm were collected for MALDI-TOF mass analysis (Shimadzu MALDI-8030) using α-cyno-4-hydroxycinnamic acid matrix (Bruker Daltonics, MA). The D-enantiomers of all peptides were purchased from GenScript and used without further modification.

### Turbidity assays

Peptide solutions (0.25–1.5 mM) were prepared in 1×PBS at pH 7.4. Aliquots (140 µL) were pipetted into a 96-well plate and absorbance at 600 nm was measured over 24 h using a Synergy HT plate reader (Biotek, USA) at RT or 37°C with or without agitation. The pH dependence of LLPS formation was assessed in 0.1 M potassium phosphate buffer (with 150 mM NaCl) at different pH values (5.8, 6.6, 7.4, 8.0) and 1 mM peptide concentration. Similarly, 0.1 M potassium phosphate buffer (pH 7.4) with varying salt concentrations (0, 150, 250, 500, 1000 mM NaCl) was used to study ionic strength effects. The absorbance was further converted to turbidity using methods previously described^66^ and data was plotted as relative turbidity.

### Optical microscopy and dynamic light scattering (DLS)

Droplets were visualized in 1×PBS using NCI Leica DMC4500 microscope in reflection mode with differential interference contrast. Images were captured with a DMC4500 camera under the control of Leica LAS X software (version 3.4.2). Coacervate size was measured using 1 mM peptide solutions (1×PBS, pH 7.4) on a ZEN 3600 Zetasizer (Malvern Instruments, UK) using DLS-Zen 0040 specialized cuvettes. The results were reported as the means of three measurements with 14 scans for each measurement.

### Transmission electron microscopy (TEM)

Peptide solutions prepared for TEM were applied directly to 200-mesh, carbon-coated copper grids for 2 min and stained with 1% uranyl formate for 1 min. Excess stain was blotted using filter paper. Brightfield images were taken with a JEOL JEM-1400 transmission electron microscope with a NANOSPRT15 camera at an accelerating voltage of 120 kV.

### Rheology testing

Viscosity of the peptide solutions (1 mM or 2 mM) was tested using an ARES 2000ex rotational rheometer (TA Instruments, New Castle, DE). Peptides were freshly solubilized, and 170 µL of each was pipetted onto the rheometer stage. A 20-mm parallel plate geometry was lowered to a gap of 200 µm. Storage modulus (G′) and loss modulus (G″) were measured as a function of angular frequency (1–10 rad/s) with 1% strain. Viscosity was measured as a function of shear rate (1–100 s^−1^). Separately, viscosity was measured using a Hagen-Poiseuille viscometer (RheoSense microVISC, San Ramon, CA) at a shear rate of 2500 s^−1^ to confirm viscosity differences at two different concentrations (1 mM and 2 mM).

### Computational simulations

All simulations were performed using GROMACS 2023 ^67,68^ and the CHARMM36m^69^ force field. The necessary force field parameters and input files were generated using AlphaFold^70^ predicted structures coupled with CHARMM-GUI ^71^. A single peptide chain was solvated in a 5-nm cubic box. Sodium and chloride ions were added to maintain electroneutrality and achieve 0.15 M NaCl. The system was energy minimized using the steepest descent algorithm and then underwent 125 ps of NVT equilibration followed by 125 ps of NPT equilibration, with 0.001 ps time steps. During these equilibration steps, the peptide atom coordinates were restrained to prevent changes. The system was further equilibrated for 125 ps in the NPT ensemble without restraints and with a 0.002 ps time step. The final equilibrated system was used for production runs with a 0.002 ps time step. Different configurations were extracted from single-chain simulations and used to build the initial multi-chain system. Ten chains, each obtained from a distinct time point or simulation of a single monomeric peptide, were incorporated into a 10-nm cubic box. Peptides were solvated and electro-neutralized, and 0.15 mM NaCl was added to achieve physiological conditions. Energy minimization was performed on the resulting system.

Next, 125 ps of NVT equilibration followed by 125 ps of NPT equilibration, with 0.001 ps time steps and no position restraints, was carried out on the energy-minimized system. The final equilibrated conformation was used for production runs with a 0.002 ps time step. For all simulations, coordinates were saved after every 1,000 steps. The V-rescale thermostat and C-rescale barostat were used for all simulations with standard temperature and pressure coupling parameters. π-π stacking interactions were calculated using a 0.5 nm distance cutoff between the centers of mass of the aromatic rings. For each pair of aromatic residues, the distance between their sidechain centers of mass was measured. If this distance was ≤ 0.5 nm and if the angle between the plane of the aromatic rings was not between 30°–60°, the pair was engaged in a π-π stacking interaction. Additionally, angles between the aromatic rings were analyzed to classify the interactions as either parallel stacking (0°–30°) or perpendicular stacking (60°–90°).

### Fourier-transform infrared (FT-IR) spectroscopy

FT-IR measurements for secondary structural analysis were conducted using 1 mM peptide solutions (1×PBS, pH 7.4) with a Bruker Alpha II FT-IR instrument equipped with a Smart Performer single-reflection ATR accessory and an Au crystal sample stage. The background spectrum and buffer spectrum were collected and subtracted from the sample using OPUS software. An average of 24 scans for each peptide were employed for each sample measurement. Data were analyzed using GRAMS/AI software (Thermo Scientific, USA). Second derivative spectra were calculated from the absorbance spectra in the Amide I region using a Savitzky-Golay filter, third order, with a nine-point window. Analyzed second derivative spectra between 1610 cm^−1^ and 1710 cm^−1^ were fit with six or seven Gaussian curves, informed by the Akaike information criterion, and the peak positions were compared to literature reports ^72–74^.

### Circular Dichroism (CD) spectroscopy

The CD spectra of the peptide solutions (0.75 mM) were recorded on a Jasco J-815 CD spectrometer. The spectra (average of three scans for each sample) were collected within the wavelength range of 215–260 nm with a bandwidth of 1.00 nm, 0.5 nm step. Solvent background was subtracted from each spectrum and data converted to mean residue ellipticity (MRE).

### Droplet coalescence assays

Experiments were conducted to visualize the dynamic process of droplet fusion by capturing a series of still images over 15 min, with each acquisition every 500 ms. Contrast and brightness were adjusted to ensure optimal visibility of the coacervates and noise cancellation was applied to reduce background interference. Frame correction aligned the images to account for shifts that occurred during capture. Additional enhancements included sharpening to visualize finer details and color correction to ensure that the visual representation closely matched the actual observation. The images were meticulously stacked to create a continuous video sequence that provided a dynamic view of the peptides’ behavior, showing the evolution of coacervates from their initial formation, through stabilization and fusion. Significant fusion events were identified and marked with arrows. Advanced tracking algorithms were employed to follow movement and changes within the coacervates, and event segmentation was used to isolate specific interactions of interest.

### Stem cell-derived cardiomyocytes and confocal microscopy

Wild type hiPSCs (WT iPSC 11) were received from the Genome Engineering and Stem Cell Center at Washington University School of Medicine. hiPSCs were cultured in mTeSR Plus medium (STEMCELL Technologies, USA) on 6-well plates coated with Matrigel (1:100, Corning) and grown at 37°C with 5% CO_2_. Cells were grown to 80–90% confluency and then passaged as clusters using Versene (Thermo Scientific, USA) into 6-well plates for hiPSC maintenance and into 24-well plates for hiPSC differentiation. Once hiPSCs reached 90% confluence, they were differentiated into cardiomyocytes using small-molecule manipulation of Wnt signaling.^75^ Non-cardiomyocytes were removed via lactate purification on days 20 and 22 before replating singularized hiPSC cardiomyocytes (hiPSC-CMs) at 40,000 cells per 35-mm glass-bottom dish.

hiPSC-CMs were treated with enantiomeric leucine or valine coacervates loaded with eGFP protein (Novus Biologicals, Catalog number: NBP2-34923; 50 μL, final concentration of 100 nM eGFP) for 30 min at 37°C. Samples were fixed with 2% paraformaldehyde (PFA), followed by three PBS washes (10 min at RT). The cell membrane was stained with wheat germ agglutinin (WGA; Fisher Scientific, Catalog number: W32466; 0.25 μL/mL) conjugated to Alexa Fluor 647 (10 min at RT), followed by three PBS washes (5 minutes at RT). Confocal micrographs of singular hiPSC-CMs were acquired using a Leica Sp8 Lightning single-photon confocal microscope equipped with five solid-state lasers (405, 488, 514, 552, and 638 nm, 12 mW each), a 63×/1.4 numerical aperture oil immersion objective, two HyD GaAsP detectors, and two high-sensitivity photomultiplier tube detectors. Images were collected as z-stacks (presented as top view, bottom view, and single plane) and by Nyquist sampling (or greater) as previously described^76^.

### BMDC cultures

All animal experiments were conducted under approved protocols by the Institutional Animal Care and Use Committee (IACUC) at Washington University in St Louis. Femoral and tibial bone marrow from C57BL/6 mice was collected and RBCs lysed using ACK buffer. The cells were plated at 5×10^6^ cells per petri dish (150 mm) and differentiated for 7–9 days in complete RPMI-1640 medium (containing 10% heat-inactivated FBS), 55 µM β-mercaptoethanol, 1 mM sodium pyruvate, 10 mM HEPES, 1× MEM non-essential amino acids, 20 ng/mL GM-CSF, 10 ng/mL IL-4, and 100 µg/mL penicillin-streptomycin (pen-strep). Non-adherent cells were collected by pipetting and washing plates gently with media and plated at required density.

### Encapsulation efficiency measurements

Stock solutions of eGFP (248 μM), FITC-H-2K^b^ antibody (BD Biosciences, Catalog number: BDB553569; 0.5 mg/mL), or BODIPY labeled ovalbumin (DQ-OVA, Catalog number: D12053, Thermo Fisher Scientific; 1 mg/mL) were solubilized in PBS and added to dry peptide powders to a final peptide concentration of 1 mM. Cargo concentrations in the stock solutions were 1 µM eGFP, 50 μg/mL FITC-H-2K^b^, and 1 mg/mL DQ-OVA. Samples were then centrifuged at 15,000 × g for 10 min, and the absorbance of the supernatant was measured for BODIPY (488 nm) and fluorescence for eGFP (λ_ex_: 395 nm and λ_em_: 510 nm) and FITC (λ_ex_: 495 nm and λ_em_: 519 nm). For small molecule dyes, similar methods were used. The encapsulation efficiency was determined as reported previously.^38^

### Cellular uptake and antigen presentation assays

Uptake of coacervates by BMDCs was measured using *in vitro* antigen presentation assays. BMDCs were plated in 96-well round-bottom plates (1×10^5^ cells/well) and maintained at 4°C or treated with 10 mM deoxy glucose and 10 mM sodium azide in PBS (ATP depletion) for 1 h prior to addition of OVA-loaded coacervates. Final OVA concentration was 1 μg/mL for time-dependent measurements and 10 μg/mL for comparison between enantiomers. Ceμls maintained at 37°C and complete RPMI media served as controls. Soluble OVA and PBS treated cells were used as controls for antigen delivery. Following treatment (2 h, 4 h, 24 h, 48 h, or 72 h), BMDCs were washed to remove extracellular coacervates and DOBW hybridoma cells (1:5, DC: DOBW) were overlaid for 16 h. The cells were then centrifuged at 300 × *g* for 5 min, and the supernatant was collected for IL-2 quantitation by ELISA according to the manufacturer protocol (Biotechne, #DY402). The amount of IL-2 produced under each condition was measured. A second antigen/hybridoma pair (Ag85B protein and BB7 hybridoma) ^77^ was used to confirm that the findings were not OVA-specific. Final antigen concentration was ∼0.5 μM and the ratio of BMDCs to BB7 cells was 1:7. Inhibition of coacervate uptake was measured by adding the inhibitors 1 h prior to coacervate treatment for 24 h followed by hybridoma overlay and IL-2 measurements. DOBW and OVA1.3 hybridoma cells were a kind gift from Dr. Clifford V. Harding (Case Western Reserve University) ^78,79^. BB7 hybridoma was a kind gift from Dr. David Canaday (Case Western Reserve University) _80._

### Cytotoxicity and transfection assays

For cytotoxicity assays, HEK293T cells or primary mouse BMDCs (5×10^4^ cells/mL in 96-well plates) were treated with peptide coacervates (100 µM) for 24 h. MTT (3-(4, 5-dimethylthiazolyl-2)-2, 5-diphenyltetrazolium bromide (Sigma-Aldrich)) solution (40 µL of 1 mg/mL stock diluted in media) was added to each well and cells were incubated further for 24 h. Media was then removed, and 150 µL DMSO was added to each well. Absorbance was measured at 570 nm using a BioTek Synergy H1 microplate reader. Untreated cells or ethanol treated cells served as controls. Cytotoxicity of all inhibitors was evaluated similarly. For transfection studies, HEK293T cells (1×10^4^ cells/mL in 24-well plates) were allowed to adhere and grow (24 h) prior to treatment with (GHGLY)_4_ or (GHGVY)_4_ coacervates loaded with eGFP DNA plasmid (Altogen Biosystems, Catalog number: 4060; 2.5 μg/mL). Cells treated with the naked plasmid or complexed with lipofectamine 2000 (3.2 µL) served as controls. GFP expression was assessed qualitatively after 96 h using a Lionheart FX automated microscope (BioTek Instruments, Inc.).

### Antigen delivery to lysosomes and inhibitor effects

DC 2.4 cells were cultured in RPMI-1640 (Catalog number: 11875093) medium supplemented with 10% FBS, 100 U/mL penicillin, 100 μg/mL streptomycin, and 1 mM sodium pyruvate. Cells were plated in 8-well chamber slides (5000 cells in 200 μL of media) and incubated for 24 h prior to addition of DQ-OVA loaded coacervates for 2 h, 6 h, or 24 h (final DQ-OVA concentration was 2.5 μg/mL). Cells treated with PBS or soluble OVA served as controls. Cells were then washed and fixed using 4% Paraformaldehyde (PFA) for 15 min at RT and permeabilized with 0.1% Triton X-100 for 5 min prior to nuclear staining with Hoechst stain (Catalog number: 62249, Thermo Fischer Scientific; 10 μg/mL). Cells were imaged (60×) on Lionheart FX automated microscope (BioTek Instruments, Inc.) with controlled exposure parameters across two channels: eGFP and DAPI. Images were captured for each time point (2, 6, and 24 h) to assess the cellular uptake and lysosomal processing of DQ-OVA. The images were deconvoluted using Gen 5 software overlaid to visualize localization. To test the effect of endocytic inhibitors on coacervate uptake, BMDCs (5×10^4^ cells/mL in 96-well plates) were pre-treated with chlorpromazine (30 µM), dynasore (50 µM), wortmannin (10 µM), IPA-3 (10 µM), or methyl β-CD (2 mM) for 1 h prior to the addition of OVA-loaded coacervates. After 6 h, cells were washed and overlaid with DOBW cells for 16 h and IL-2 production was assessed by ELISA as described above. The effect of MHC I inhibitors bortezomib (10 nM), brefeldin A (17.8 µM) and lactacystin (50 µM) or MHC II inhibitors NH_4_Cl (20 mM), bafilomycin (500 nM), and 3-methyl adenine (10 µM) was tested similarly, using appropriate hybridomas for MHC II (DOBW) or MHC I (OVA 1.3) presentation.

### T cell proliferation and functional cytokine production

Cultured mouse BMDCs (7-9 days) were plated in 96-well plates at a concentration of 5×10^4^ cells per well and treated with OVA loaded coacervates for 24 h. Spleens and lymph nodes were harvested from OT-I/OT-II transgenic mice and single-cell suspensions were prepared by mechanical dissociation in EasySep Buffer (1× DPBS, 2% FBS, 1 mM EDTA). Cells were centrifuged at 300 × *g* for 10 min, counted using trypan blue exclusion staining, and resuspended at 1×10^8^ cells/mL in EasySep Buffer. CD8^+^/CD4^+^T cells (for OT-I/OT-II mice) were isolated from this suspension using the EasySep Mouse CD8^+^/CD4^+^ T Cell Isolation kit (STEMCELL Technologies; Cat#19852 for CD4^+^T; Cat#19853 for CD8^+^T) *via* negative selection using the Big Easy EasySep magnet (STEMCELL Technologies; Cat#18001) following the manufacturer’s instructions. Isolated CD8^+^/CD4^+^T cells were washed twice with 1× PBS, counted again, and resuspended at 5×10^6^ cells/mL in 1× PBS containing 2 µM CellTrace Violet (CTV, Thermo Fisher, USA Cat#C34571). Cells were incubated at 37°C (protected from light) for 20 min, after which 5× staining volume of EasySep Buffer was added for another 5 min of incubation. The cells were counted again, centrifuged at 300 *g* for 5 min, resuspended in RPMI-1640 (Gibco 31800-022) with 10% FBS (R and D Systems S11150H), 10 mM HEPES (Sigma H0887), 1× MEM NEAA (Gibco 11140-050), 1 mM sodium pyruvate (Gibco 11360-070), 1× pen-strep glutamine (Gibco 10378-016), and 55 µM β-mercaptoethanol (Gibco 21985-023) added fresh, and then plated onto treated BMDCs. The plates were incubated for ∼66 h, after which supernatants were collected for Mouse Adaptive Immune CodePlex Secretome analyses (Bruker Cellular Analysis, Inc), and cells were collected for flow cytometry studies.

### Flow cytometry

Total cells from spleen and lymph nodes of OT-I/OT-II mice were washed with FACS wash buffer (FWB; 1× PBS with 10% FBS) and transferred into a round-bottom 96-well plate for staining. Cells were washed again with 1×PBS and then resuspended in 100 µl of PBS with Zombie NIR (for OT-I; Biolegend Cat#423105) and 7-AAD (for OT-II; Biolegend Cat#420404) and anti-mouse CD16/32 (Biolegend Cat#101302) antibody. After a 30 min incubation, staining was quenched. The cells were centrifuged, washed, and resuspended in 100 µl FWB with APC anti-mouse CD8 antibody (Biolegend Cat#17-0081-82) and APC/Cy7 (Biolegend Cat#100413) anti-mouse CD4 antibody for OT-I and OT-II mice, respectively. After 30 min, staining was quenched, cells washed twice, and resuspended in 200 µl FWB for acquisition. Single-stained cells or UltraComp eBeads (Invitrogen Cat#01-3333-42) stained with Zombie NIR, CTV, or CD8 and 7-AAD, CTV, or CD4 APC/Cy7 were used as compensation controls for OT-I and OT-II mice, respectively. Samples were acquired using an Agilent NovoCyte 3000 flow cytometer. Data were analyzed with FlowJo version 10. Proliferation was quantified as the percentage of daughter cells out of single, live, CD4^+^ or CD8^+^T cells, gated as having a lower CTV fluorescence intensity than the peak of highest fluorescence (undivided).

### Statistical analyses

Statistical analysis was performed in GraphPad Prism. Data are expressed as mean ± S.E.M. and statistical analysis was performed using a one-way or two-way ANOVA with Tukey/Sidak’s/Dunnett’s multiple comparison test. *p ≤ 0.05, **p ≤ 0.01, ***p ≤ 0.001, ****p ≤ 0.0001.

## ACKNOWLEDGEMENTS

Research funding was provided through the Washington University McKelvey School of Engineering, Department of Biomedical Engineering Commitment Funds (12–360–94361 J) and the National Institute of Health, National Institute of Allergy and Infectious Diseases (R01 AI168918) and National Science Foundation CAREER award to Dr. Jai Rudra. Jagannath Mondal acknowledges the support of the Department of Atomic Energy, Government of India, under Project Identification No. RTI 4007.

