## Supporting Information for "Enantiomeric histidine-rich peptide coacervates enhance antigen delivery to T cells"

**Figure S1**. MALDI-TOF spectra of L-form (GHGXY)_4_ peptides	S4

**Figure S2** MALDI-TOF spectra of D-form (GHGXY)_4_ peptides	S5

**Figure S3.** HPLC analysis of L-form (GHGXY)_4_ peptides	S6

**Figure S4**. HPLC analysis of D-form (GHGXY)_4_ peptides	S7

**Figure S5** LLPS of (GHGXY)_4_ peptides in water	S8

**Figure S6** LLPS of (GHGXY)_4_ peptides in 150 mM NaCl	S9

**Figure S7** TEM images of (GHGXY)_4_ peptides	S10

**Figure S8** Determination of C_crit_ for LLPS	S11

**Table S1**: C_crit_ for L-form and D-form peptides	S11

**Figure S9** Kinetics of LLPS at A_600_ without shaking	S12

**Figure S10** Kinetics of LLPS at A_350_	S13

**Figure S11**. Optical images of droplets at different NaCl concentration	S14

**Figure S12**. Optical images of droplets at different pH	S15

**Figure S13**. Effect of temperature (37°C) on LLPS kinetics	S16

**Figure S14**. FT-IR spectra of L-form and D-form peptides	S17

**Figure S15**. Storage and loss modulus of (GHGXY)_4_ peptide solutions	S18

**Figure S16**. Parallel and perpendicular π-stacking interactions from all-atom simulations	S19

**Figure S17**. Microscopy images of GFP loaded (GHGXY)_4_ droplets	S20

**Figure S18**. Encapsulation efficiency of (GHGXY)_4_ droplets	S21

**Figure S19**. 3-D Top view images of GFP loaded coacervates in hiPSCs	S22

**Figure S20**. Representative coalescence images of (GHGLY)_4_ droplets	S23

**Figure S21**. Controls depicting no fluorescence in the absence of coacervates	S24

**Figure S22.** Delivery of eGFP plasmid DNA in HEK293T cells.	S25

**Figure S23**. Cytotoxicity of coacervates in HEK293T cell line	S26

**Figure S24.** Cytotoxicity of coacervates in primary murine BMDCs	S27

**Figure S25.** Cytokine produciton by BMDCs stimulated coacervates	S28

**Figure S26**. DQ-OVA fluorescence BMDCs at various time points	S29

**Figure S27.** Cytotoxicity of inhibitors used in the study	S30

**Figure S28**. DQ-OVA fluorescence intensity in the presence of inhibitors	S31

**Figure S29**. Antigen presentation using Ag85B protein and BB7 hybridoma cells	S32

**Figure S30**. Flow data for transgenic CD4^+^T cells	S33

**Figure S31**. Cytokine production by transgenic CD4^+^T cells	S34

**Figure S32**. Flow data for transgenic CD8^+^T cells	S35

**Figure S33**. Cytokine production by transgenic CD8^+^T cells	S36

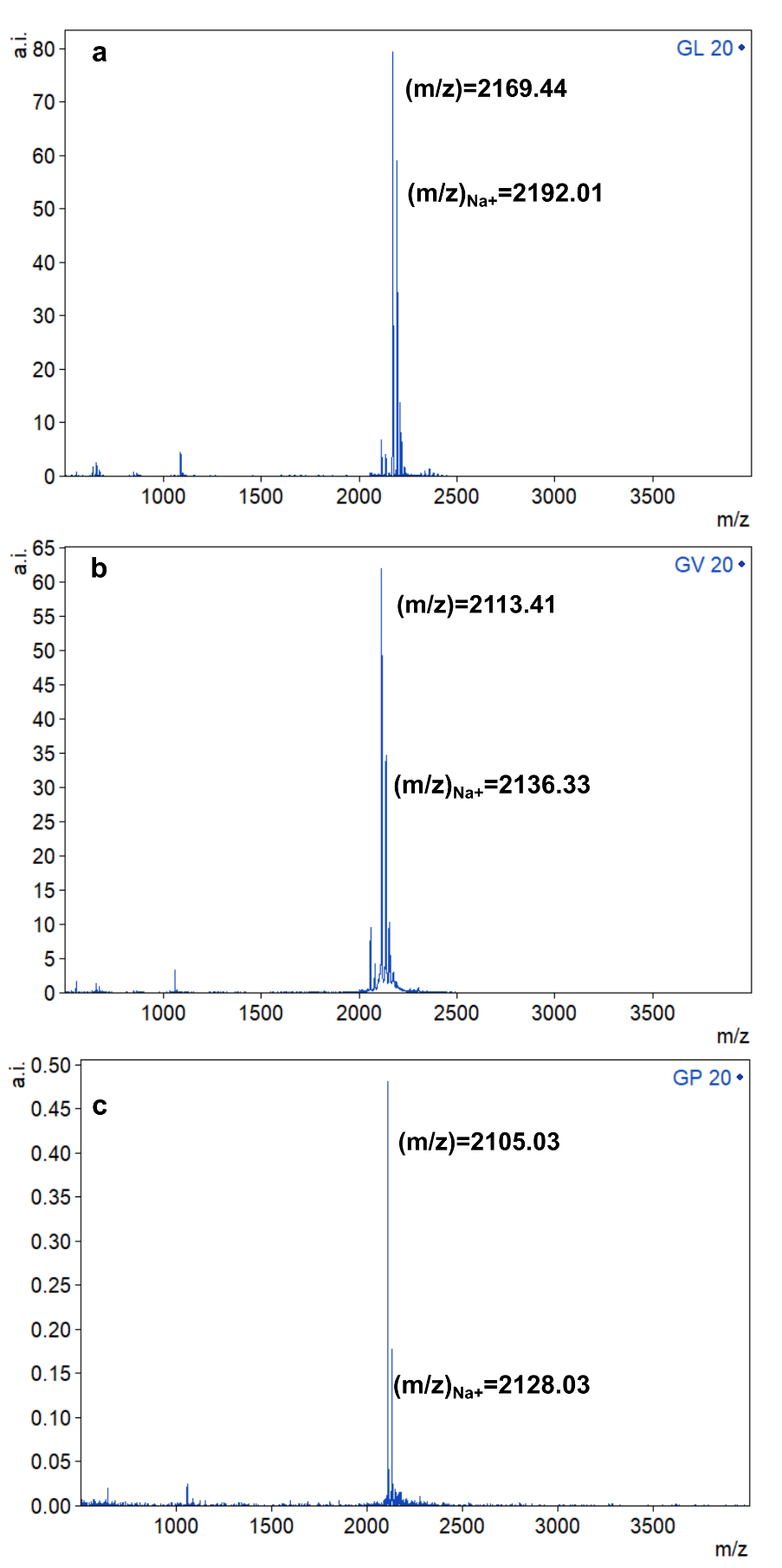

**Figure S1:** MALDI-TOF spectra of L-form (a) (GHGLY)_4_, (b) (GHGVY)_4_, and (c) (GHGPY)_4_.

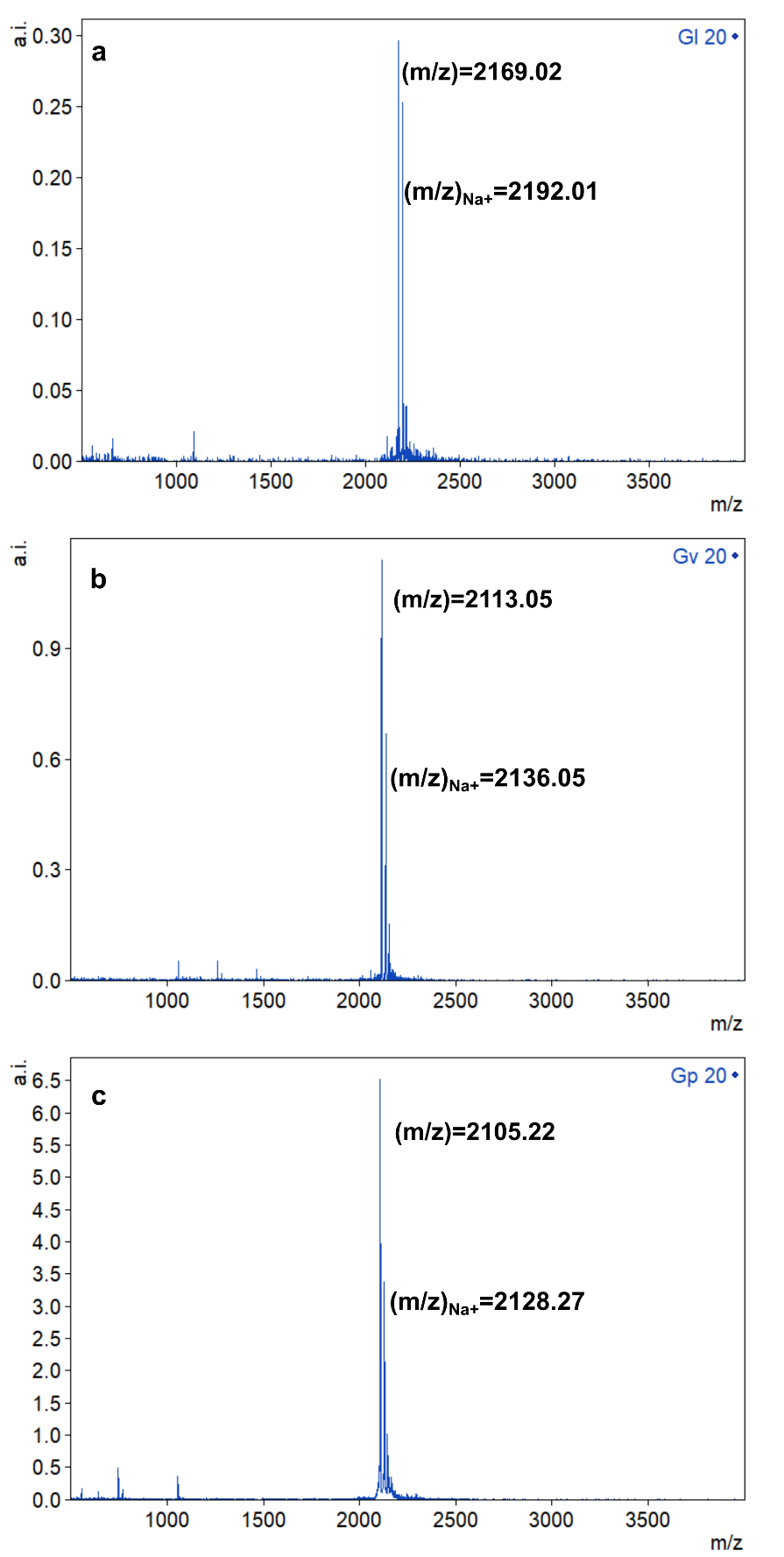
**Figure S2:** MALDI-TOF spectra of D-form (a) (GHGLY)_4_, (b) (GHGVY)_4_, and (c) (GHGPY)_4_.

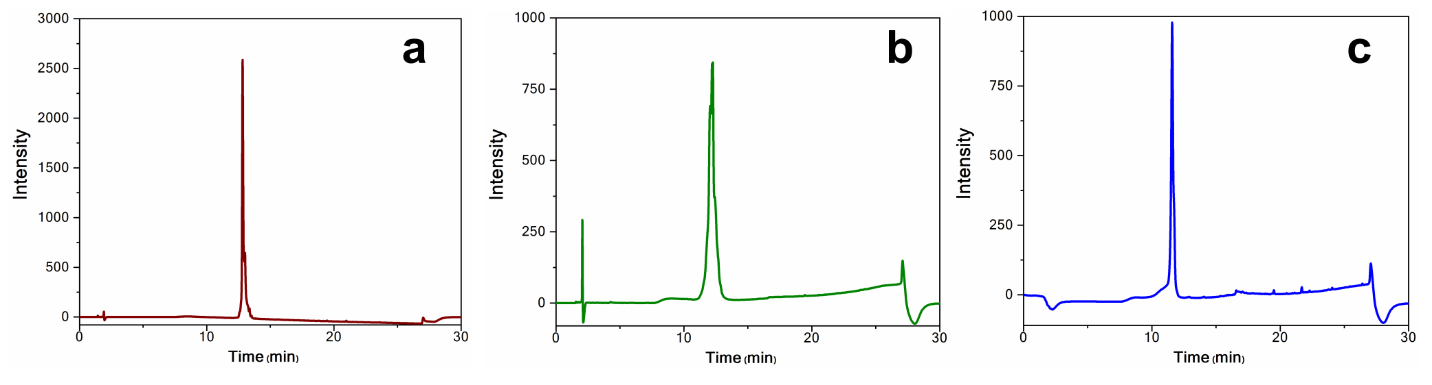
**Figure S3.** HPLC chromatograms of purified L-form (a) (GHGLY)_4_, (b) (GHGVY)_4_, and (c) (GHGPY)_4_.

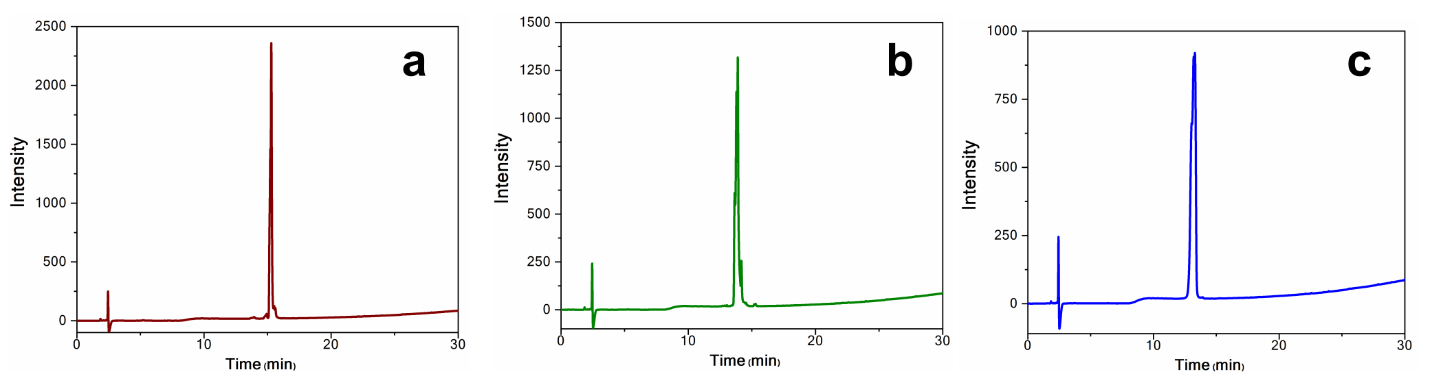

**Figure S4.** HPLC chromatograms of purified D-form (a) (GHGLY)_4_, (b) (GHGVY)_4_, and (c) (GHGPY)_4_.

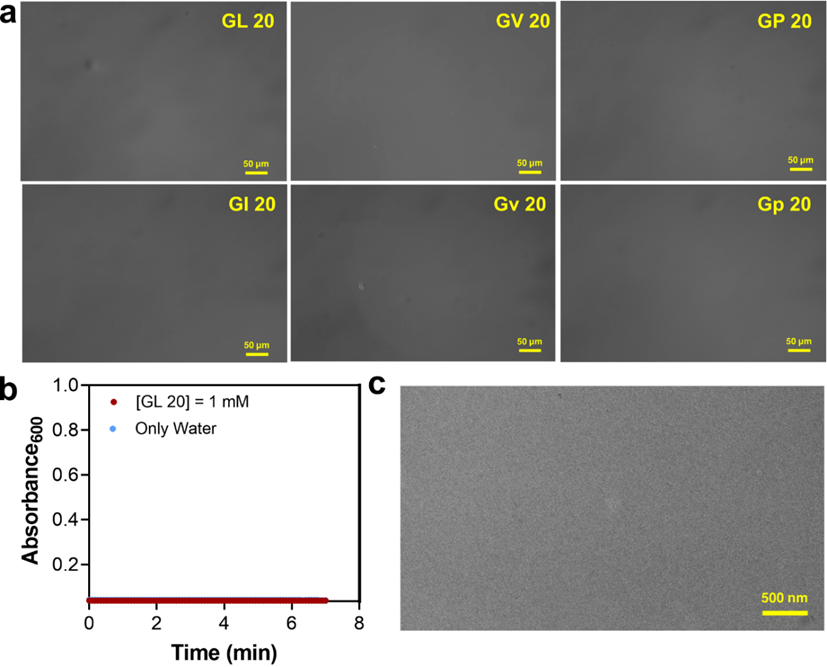

**Figure S5.** TEM images of (a) (GHGLY)_4_, (GHGVY)_4_, and (GHGPY)_4_ and their enantiomers, (b) kinetics of LLPS by 1 mM (GHGLY)_4_ in water, and (c) TEM image of 1 mM (GHGLY)_4_ in water.

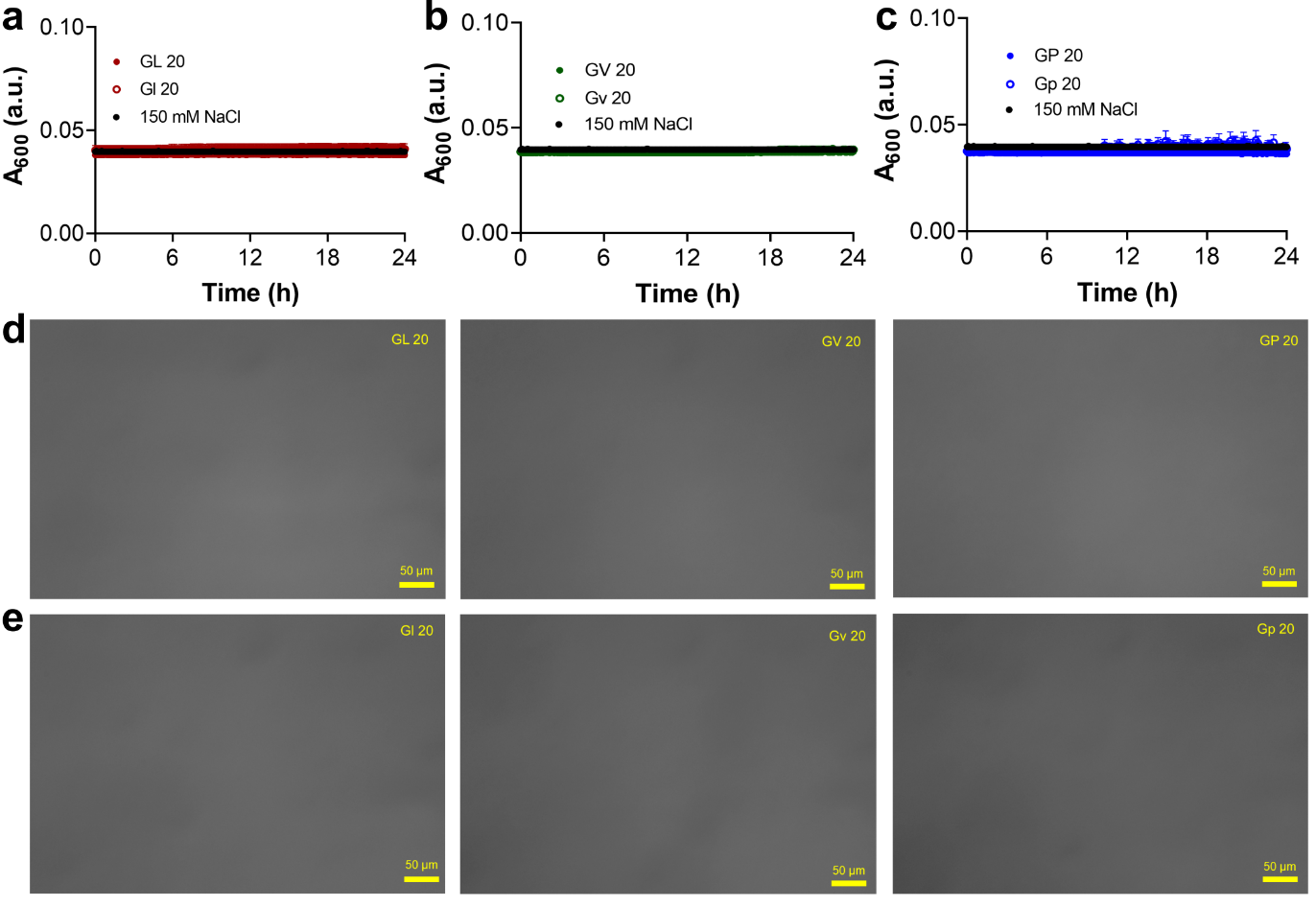

**Figure S6.** Turbidity measurements of (a) (GHGLY)_4_, (b) (GHGVY)_4_, (c) (GHGPY)_4_ peptides and their enantiomers (1 mM, in 150 mM NaCl) and corresponding optical micrographs of (d) L-form peptides and (e) D-form peptides.

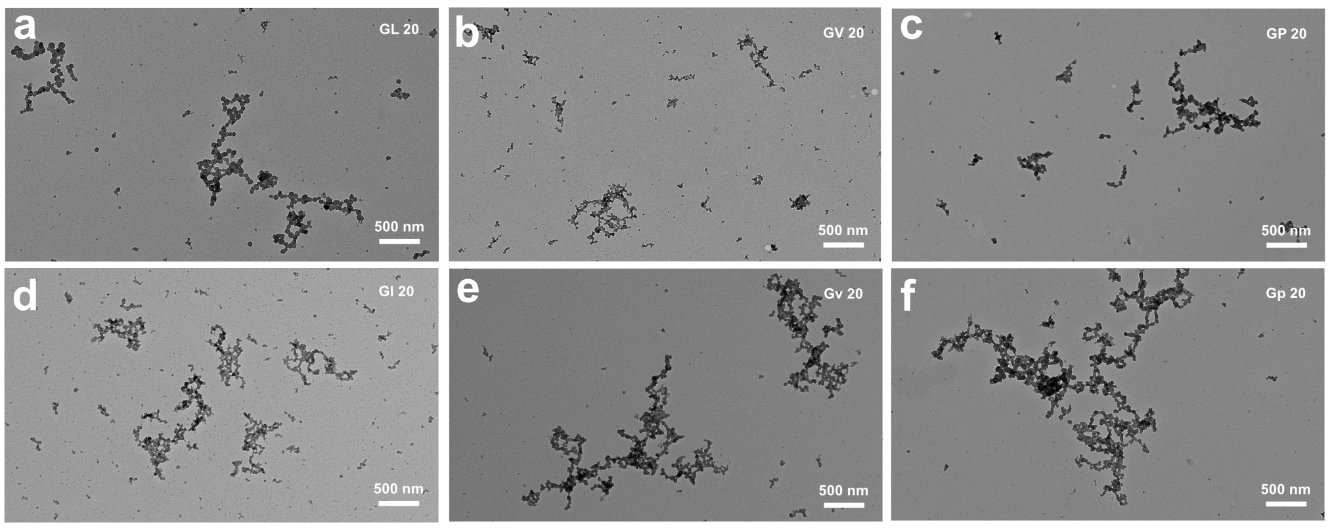

**Figure S7.** TEM images of droplets formed by (a) (GHGLY)_4_, (b) (GHGVY)_4_, (c) (GHGPY)_4_ and (d-f) their corresponding enantiomers (1 mM, 1× PBS, pH 7.4).

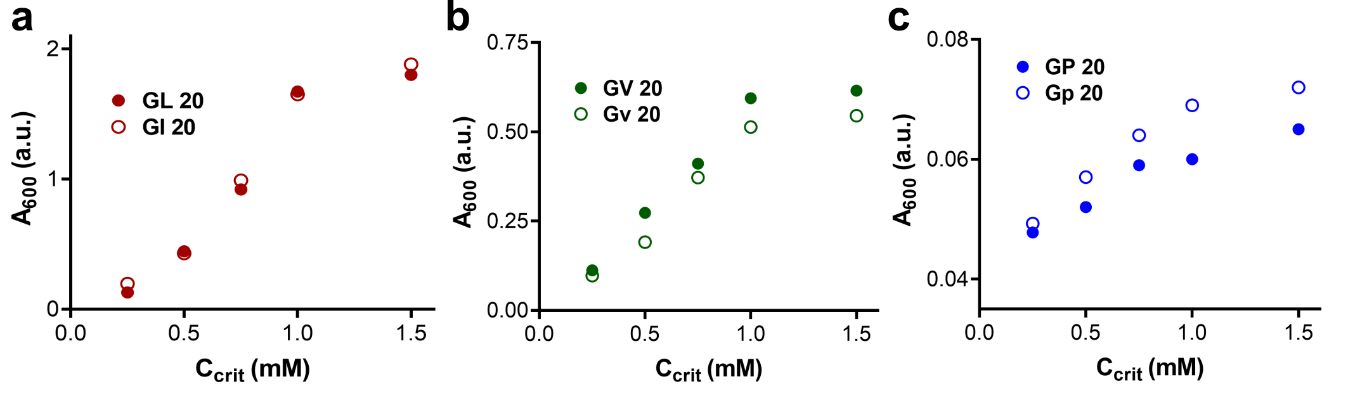

**Figure S8.** Determination of C_crit_ for (a) (GHGLY)_4_, (b) (GHGVY)_4_, (c) (GHGPY)_4_ and (d-f) their corresponding enantiomers (1 mM, 1× PBS, pH 7.4).

**Table S1. Measured** C_crit_ values for (GHGXY)_4_ peptides used in this study.

| **Peptide** | **C_crit_ (mM)** |
| --- | --- |
| GL 20/Gl 20 | 0.5 mM |
| GV 20/Gv 20 | 0.75 mM |
| GP 20/Gp 20 | 1 mM |

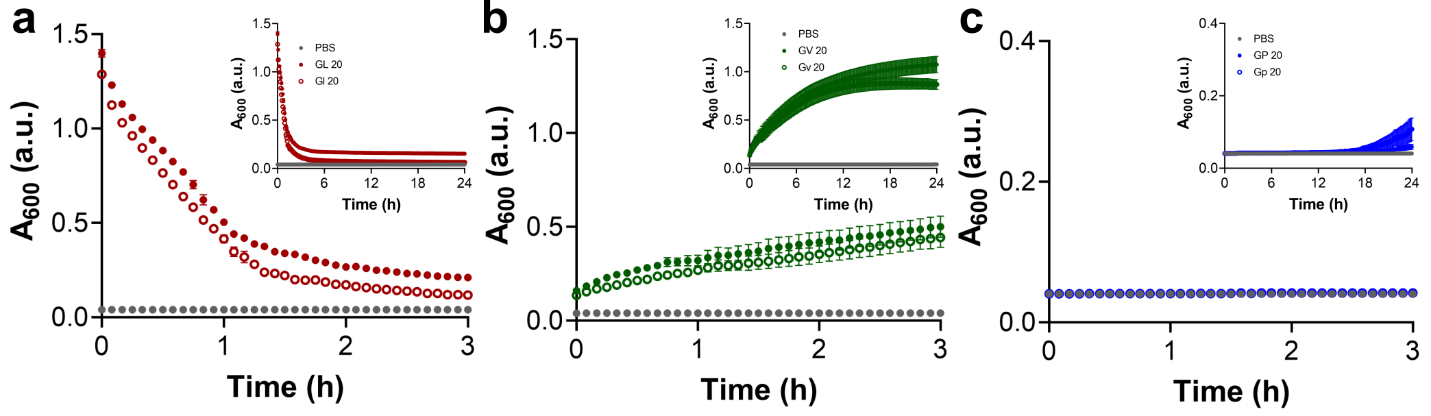

**Figure S9.** Turbidity measurements at 600 nm for (a) (GHGLY)_4_, (b) (GHGVY)_4_, (c) (GHGPY)_4_ and their corresponding enantiomers (1 mM, 1×PBS, pH 7.4) without shaking. Inset shows measurements up to 24 h.

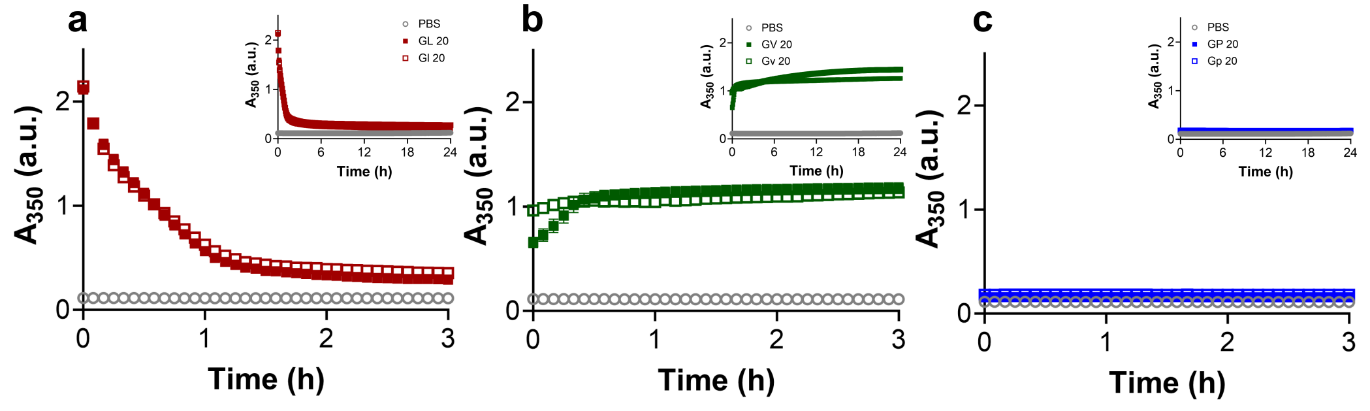

**Figure S10.** Turbidity measurements at 350 nm for (a) (GHGLY)_4_, (b) (GHGVY)_4_, (c) (GHGPY)_4_ and their corresponding enantiomers (1 mM, 1× PBS, pH 7.4). Inset shows measurements up to 24 h.

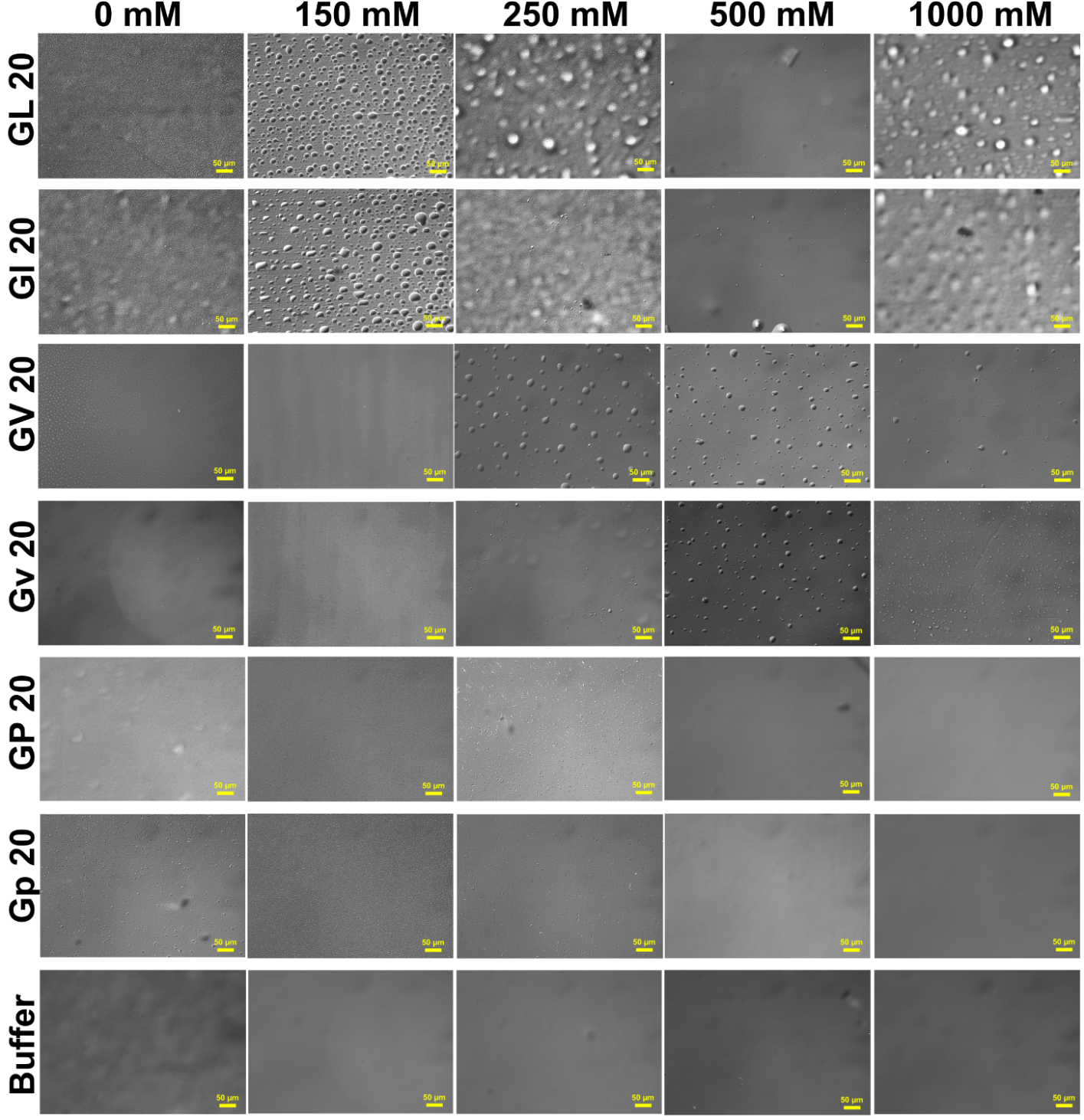

**Figure S11.** Optical images of droplets formed by L-form and D-form (GHGXY)_4_ variants at different salt concentrations (0-1000 mM) in 0.1 M phosphate buffer (pH 7.4).

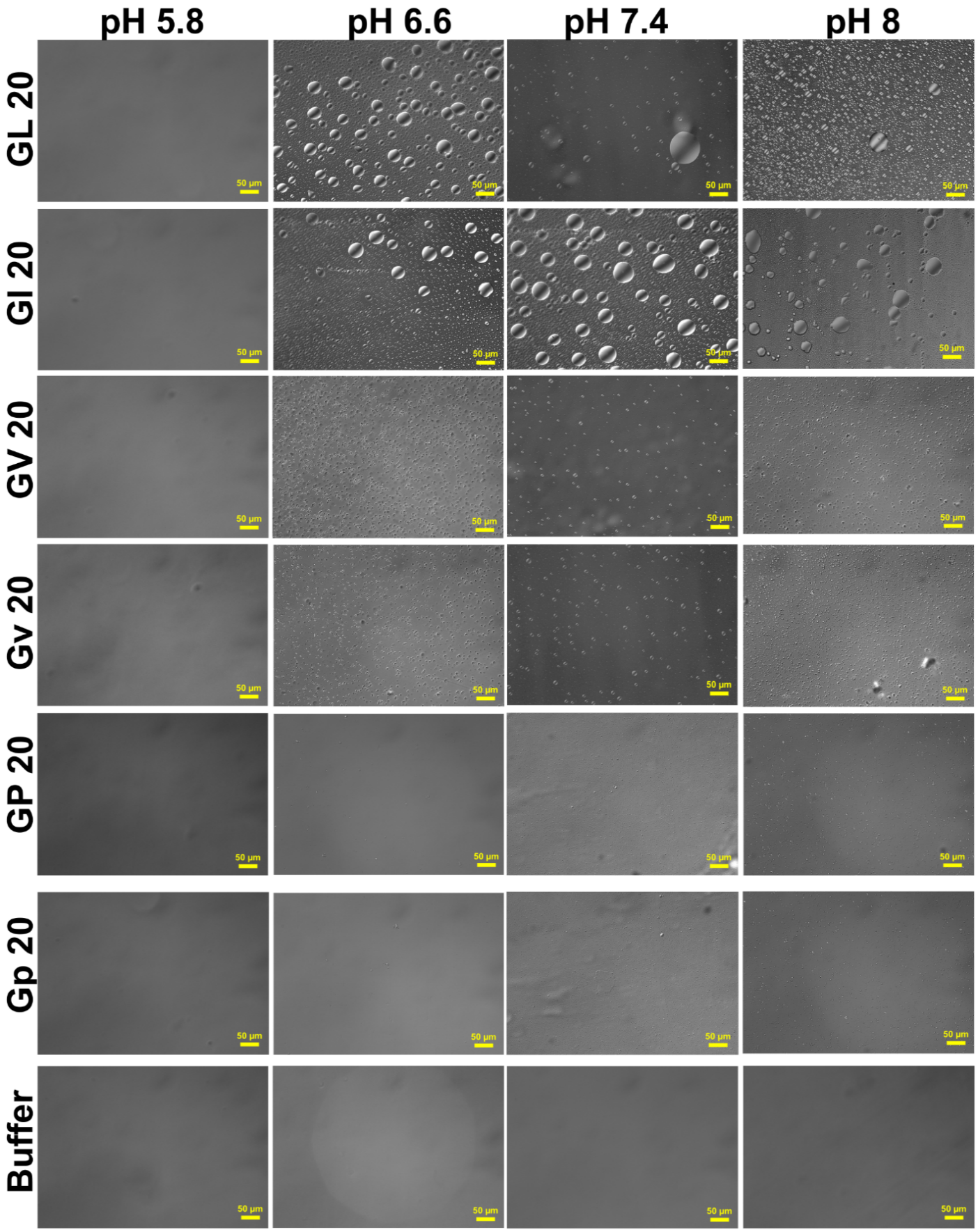

**Figure S12.** Optical images of droplets formed by L-form and D-form of (GHGXY)_4_ variants with varying pH of 0.1 M phosphate buffer.

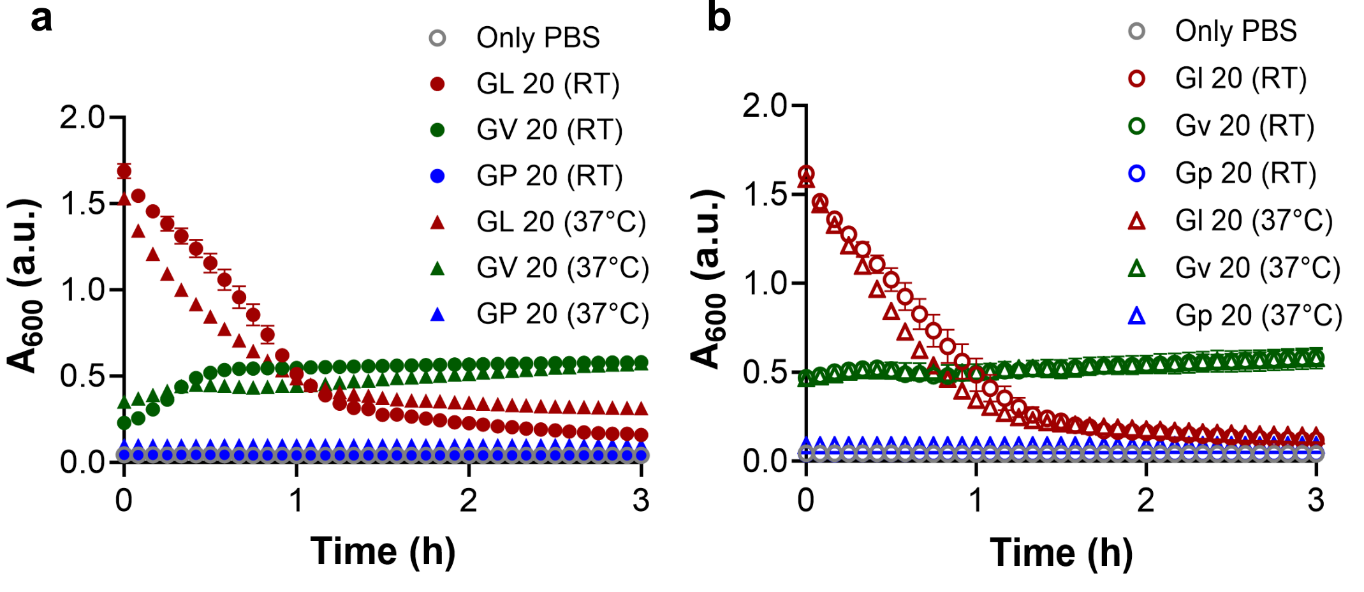

**Figure S13.** Turbidity measurements of (a) L-form (GHGXY)_4_ variants and (b) D-form (GHGXY)_4_ variants at 37°C (circles) and RT (triangles) in 1×PBS, pH 7.4.

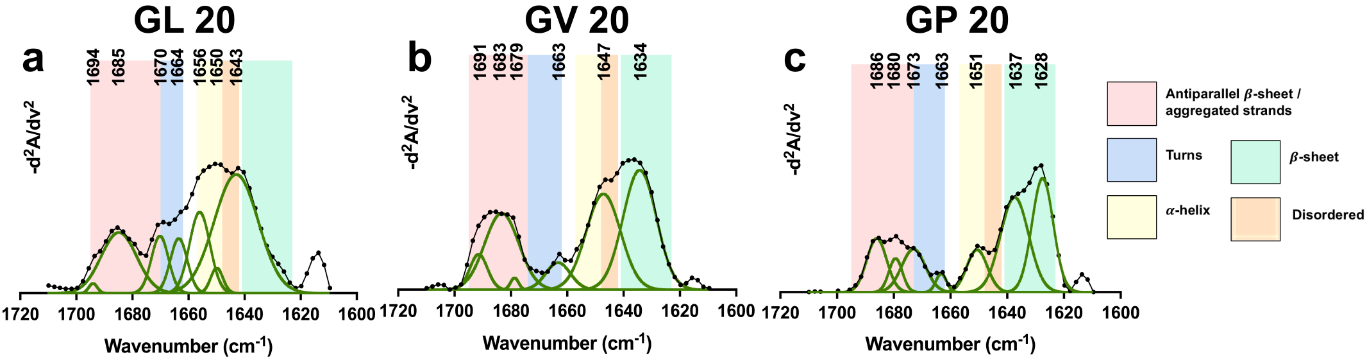

**Figure S14.** Second-derivative FT-IR spectra of (a) (GHGLY)_4_, (b) (GHGVY)_4_, and (c) (GHGPY)_4_ peptide coacervates in 1×PBS (pH 7.4).

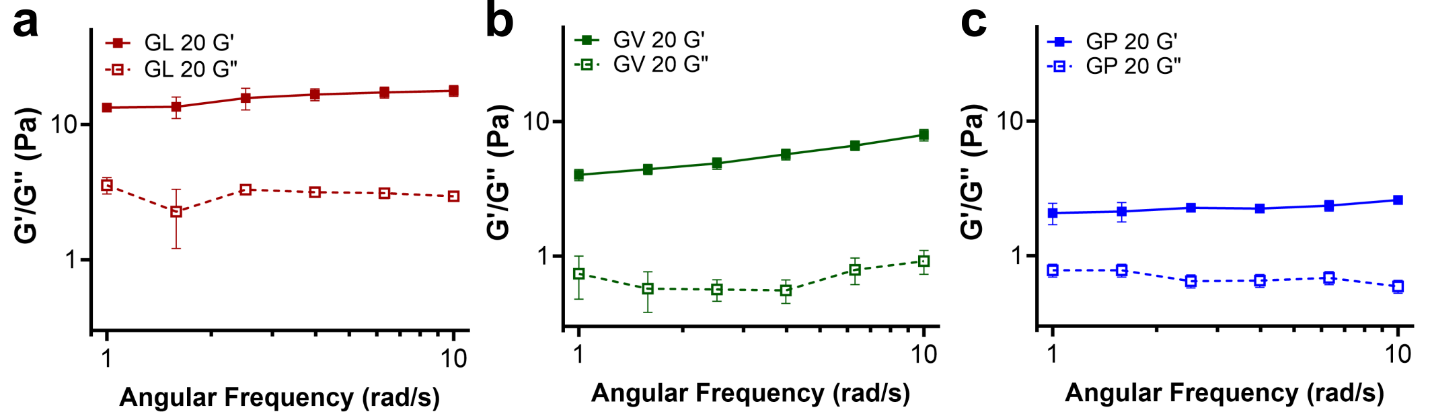

**Figure S15.** Storage (*G*′) and loss modulus (*G*″) of (a) (GHGLY)_4_, (b) (GHGVY)_4_, and (c) (GHGPY)_4_ coacervates (1 mM, 1×PBS, pH 7.4).

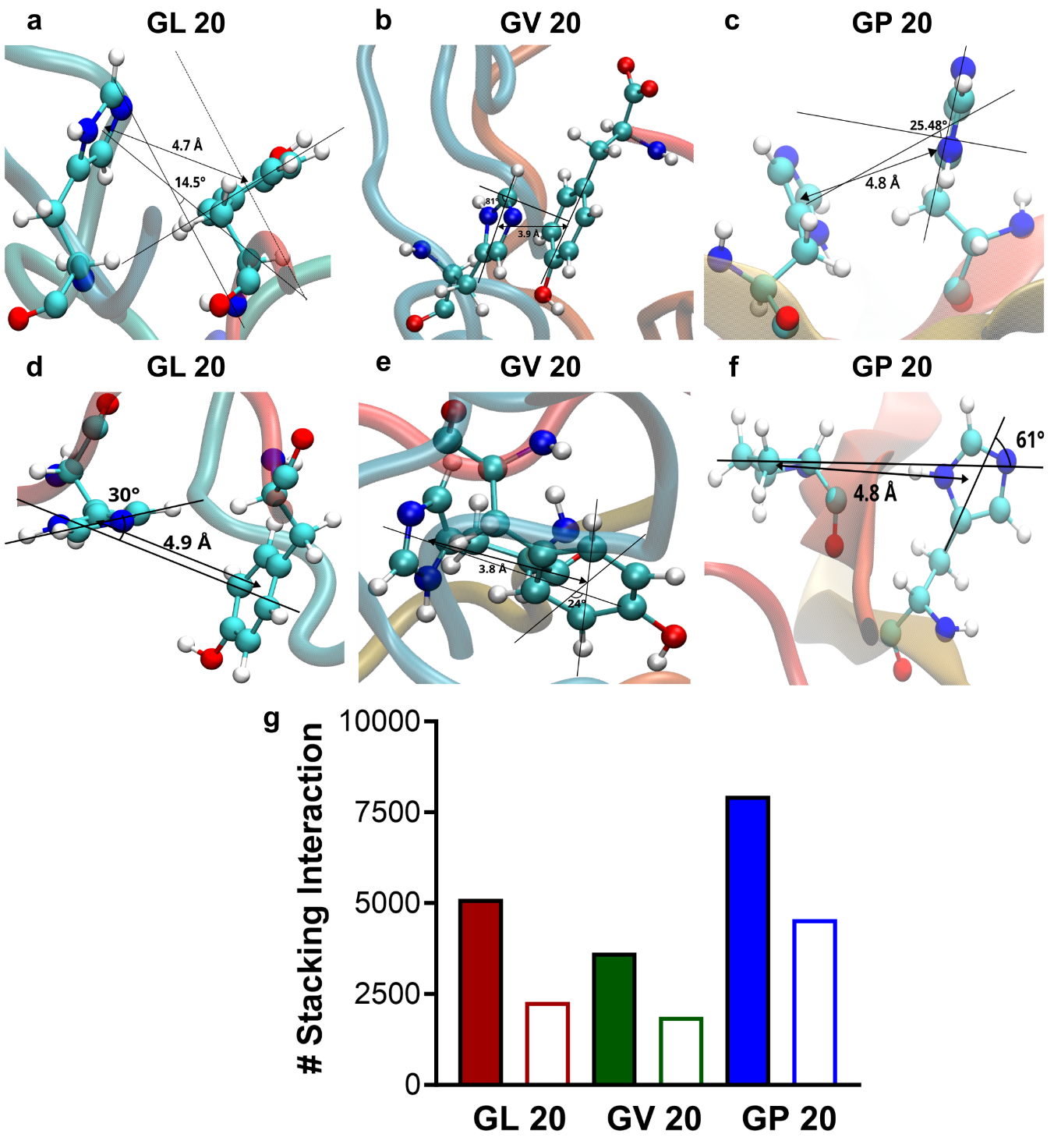

**Figure S16.** Parallel π-stacking interactions for (a) (GHGLY)_4_, (b) (GHGVY)_4_, and (c) (GHGPY)_4_ peptides. Perpendicular π-stacking interactions for (d) (GHGLY)_4_, (e) (GHGVY)_4_, and (f) (GHGPY)_4_ peptides. (g) Number of stacking parallel (solid bars) and perpendicular (clear bars) interactions occurred during the last 500 ns of simulation trajectories for all three peptides.

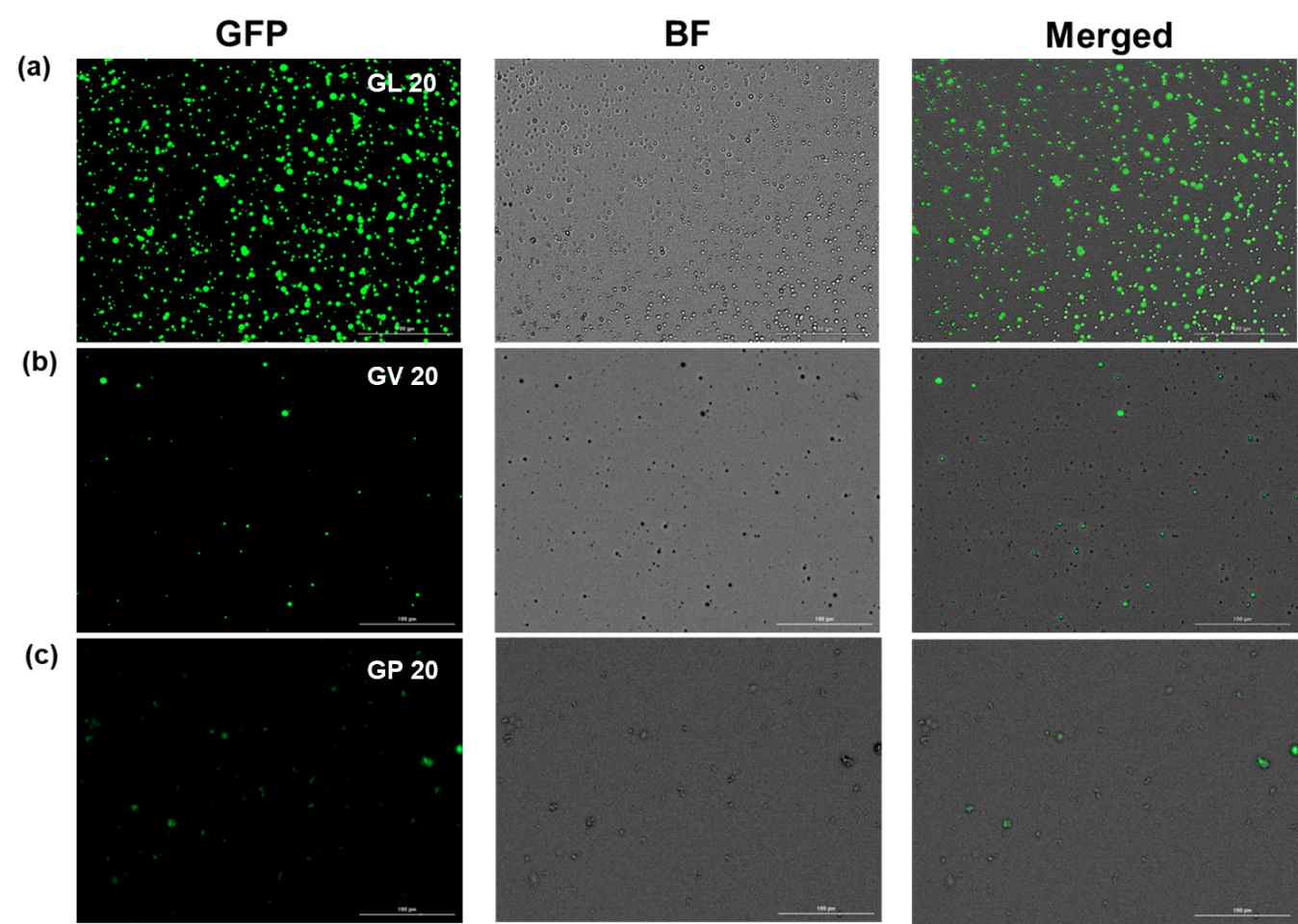

**Figure S17.** Fluorescence, bright field, and merged microscopy images of eGFP loaded (a) (GHGLY)_4_, (b) (GHGVY)_4_, and (c) (GHGPY)_4_ coacervates. Left to right panels represent GFP, DIC and merged channels. Scale bar = 100 µm.

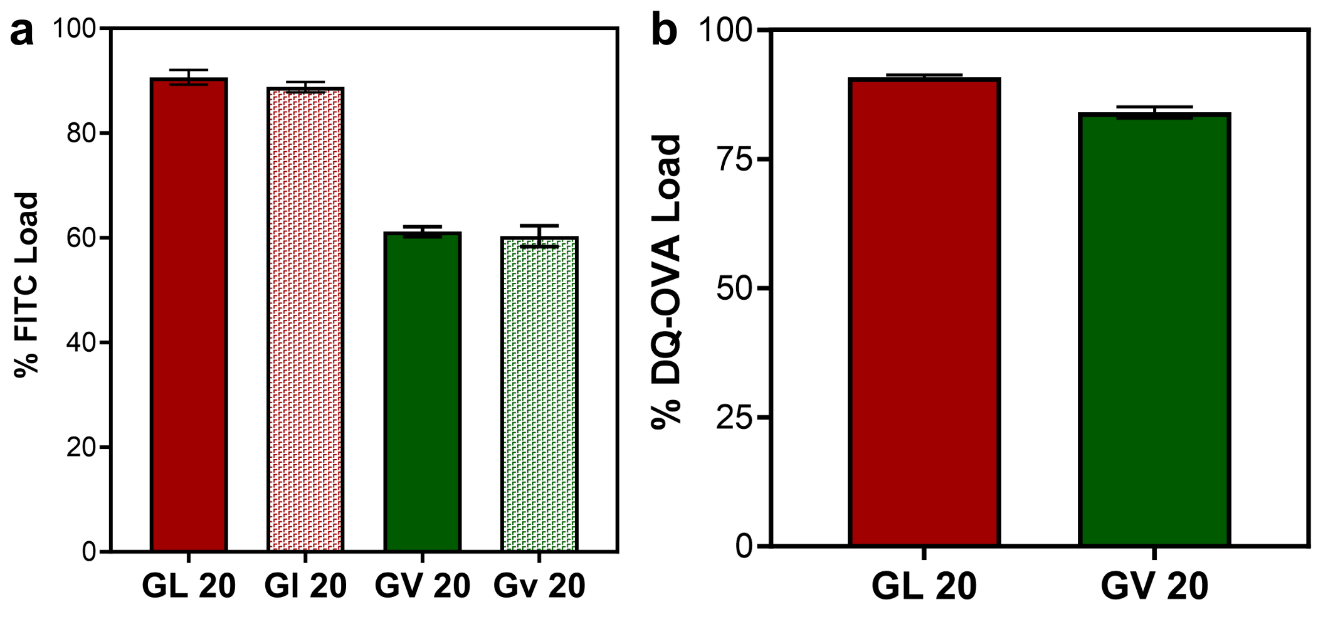

**Figure S18.** Encapsulation efficiency of (a) FITC-H-2K^b^ antibody by leucine or valine coacervates and their enantiomers and (b) model antigen DQ-OVA load by leucine and valine droplets.

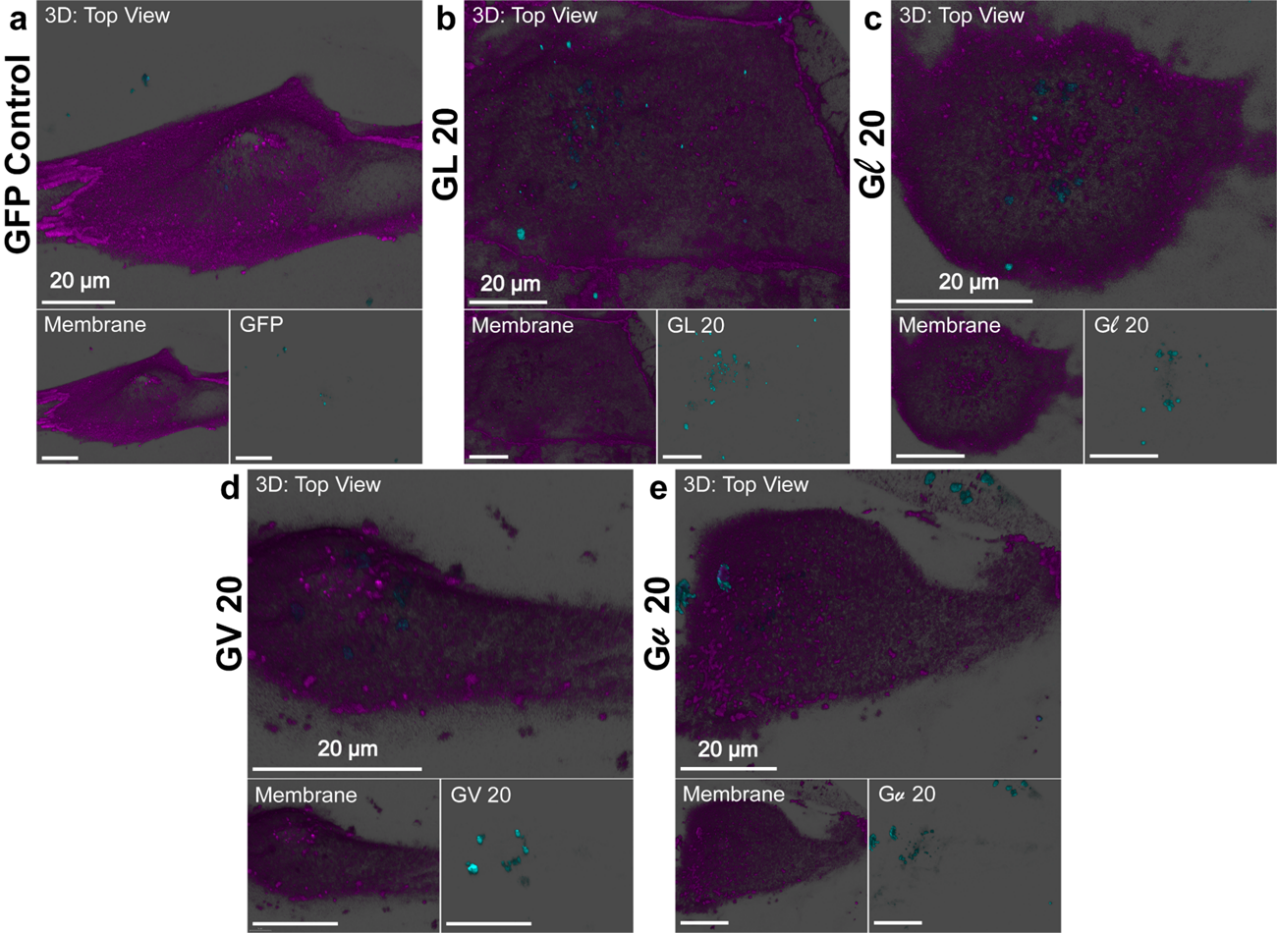

**Figure S19.** 3-D Top view of eGFP loaded coacervates in hiPSC-CM cultures. (a) GFP Control, (b, c) (GHGLY)_4_ and enantiomer (d, e) (GHGVY)_4_, and enantiomer.

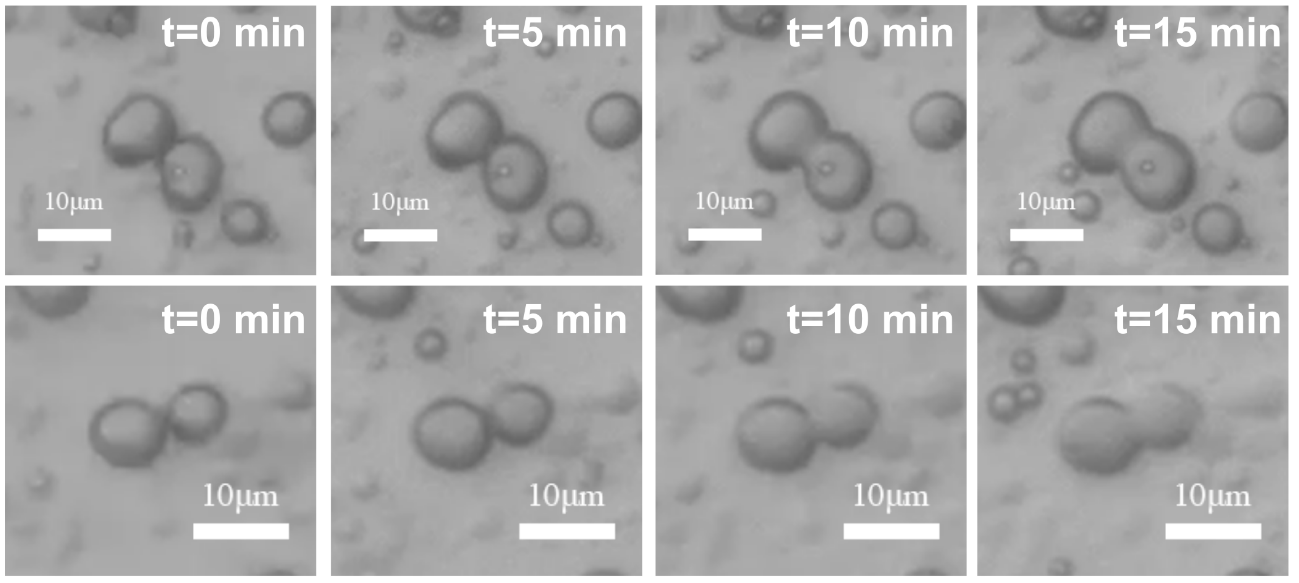

**Figure S20.** Fusion of (GHGLY)_4_ droplets shown in different time-lapse frames. Top and bottom panels show two different events.

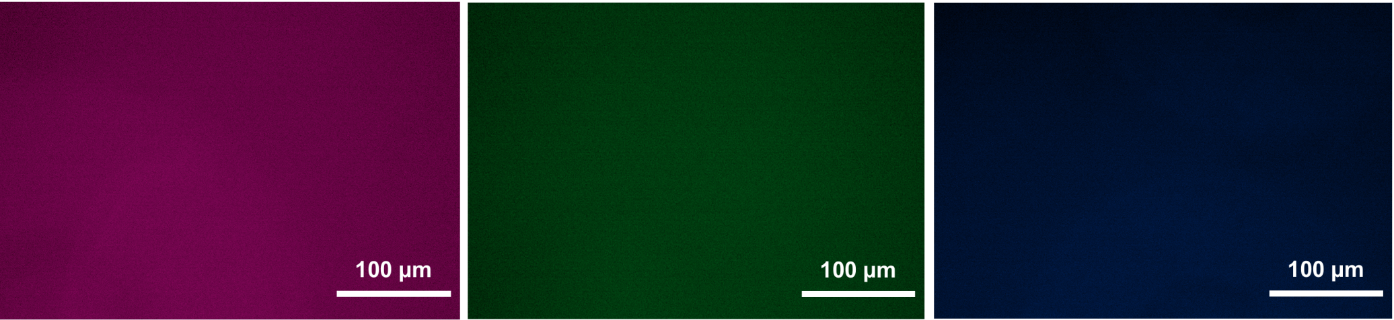

**Figure S21.** Control microcopy images showing no fluorescent puncta in the absence of coacervates (Left-Red, Middle-Green, Right-Blue).

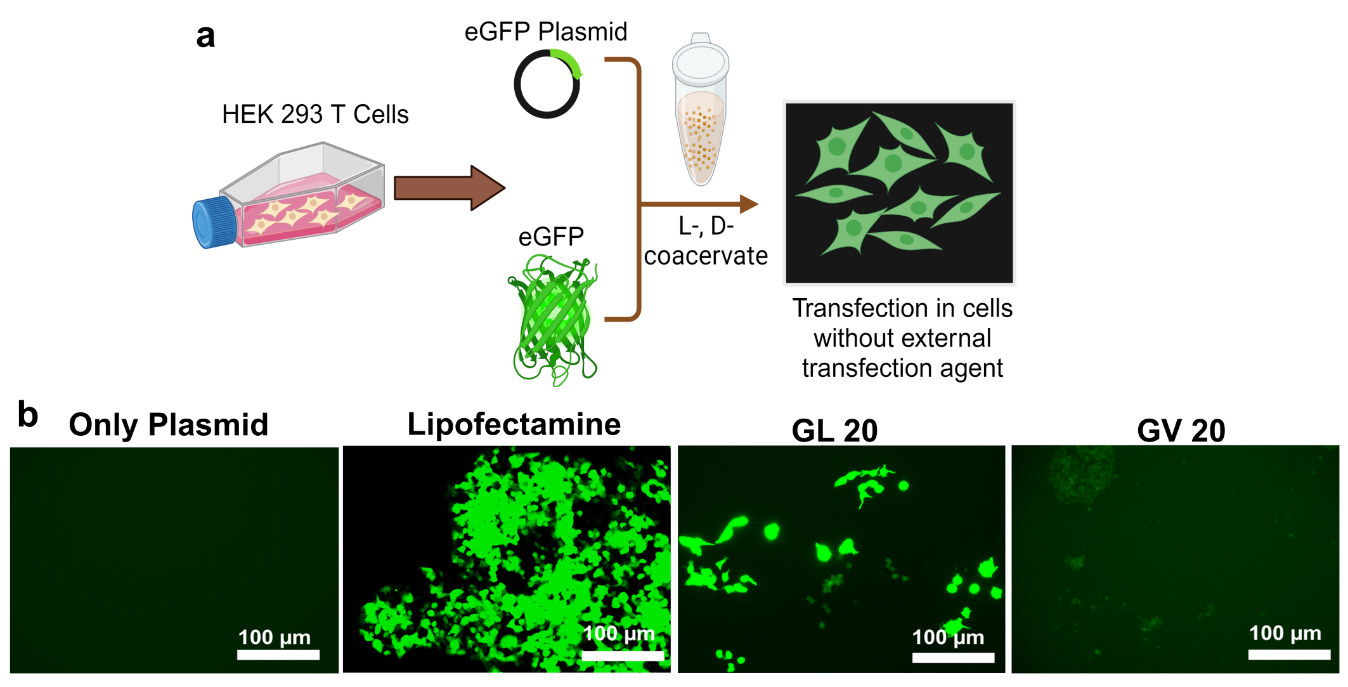

**Figure S22.** (a) Schematic showing the delivery of eGFP DNA plasmid. (b) Fluorescence micrographs of HEK293T cells transfected with (b) naked plasmid, lipofectamine-plasmid complexes or plasmid loaded in (GHGLY)_4_ or (GHGVY)_4_ coacervates. Images were taken 96 h after transfection.

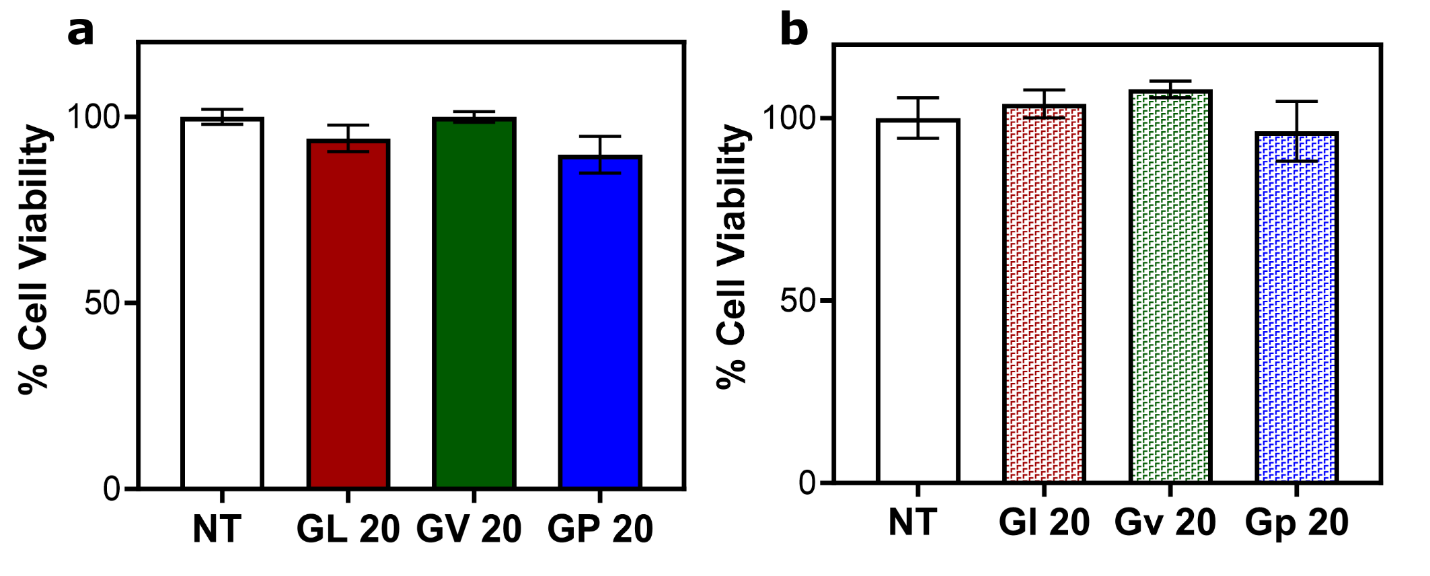

**Figure S23.** Viability of HEK-293T cells following treatment with (a) L-form or (b) D-form (GHGXY)_4_ coacervates (100 μM) for 24 h .

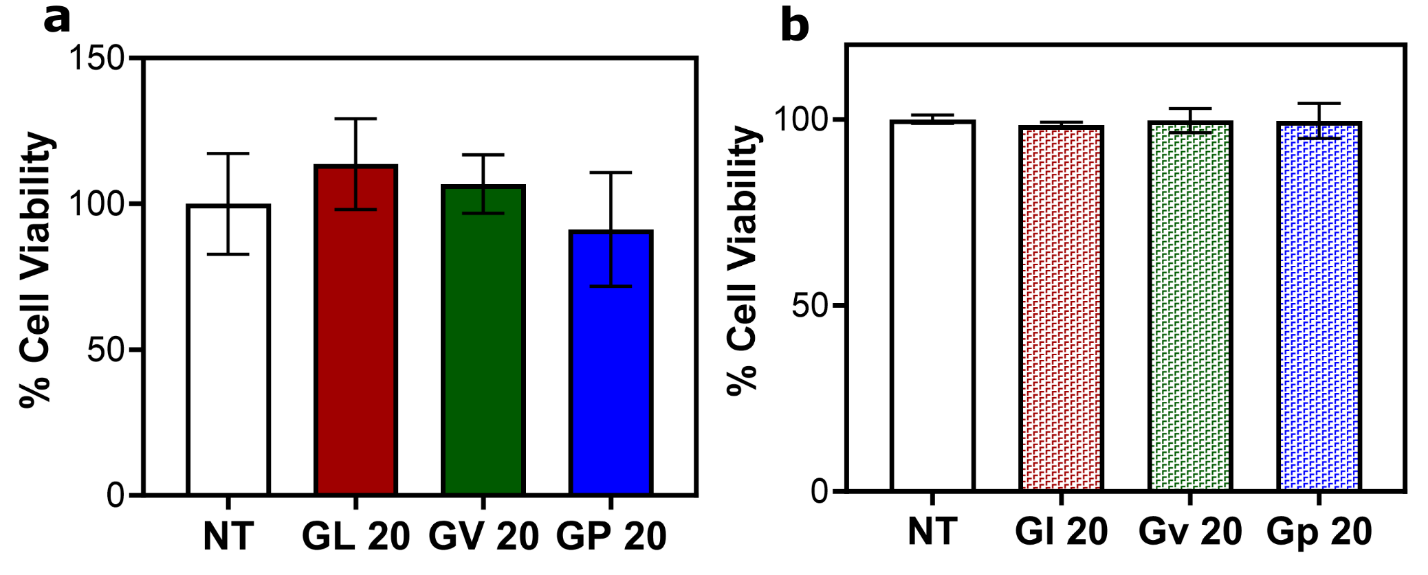

**Figure S24.** Viability of primary murine BMDCs following treatment with (a) L-form or (b) D-form (GHGXY)_4_ coacervates (100 μM) for 24 h .

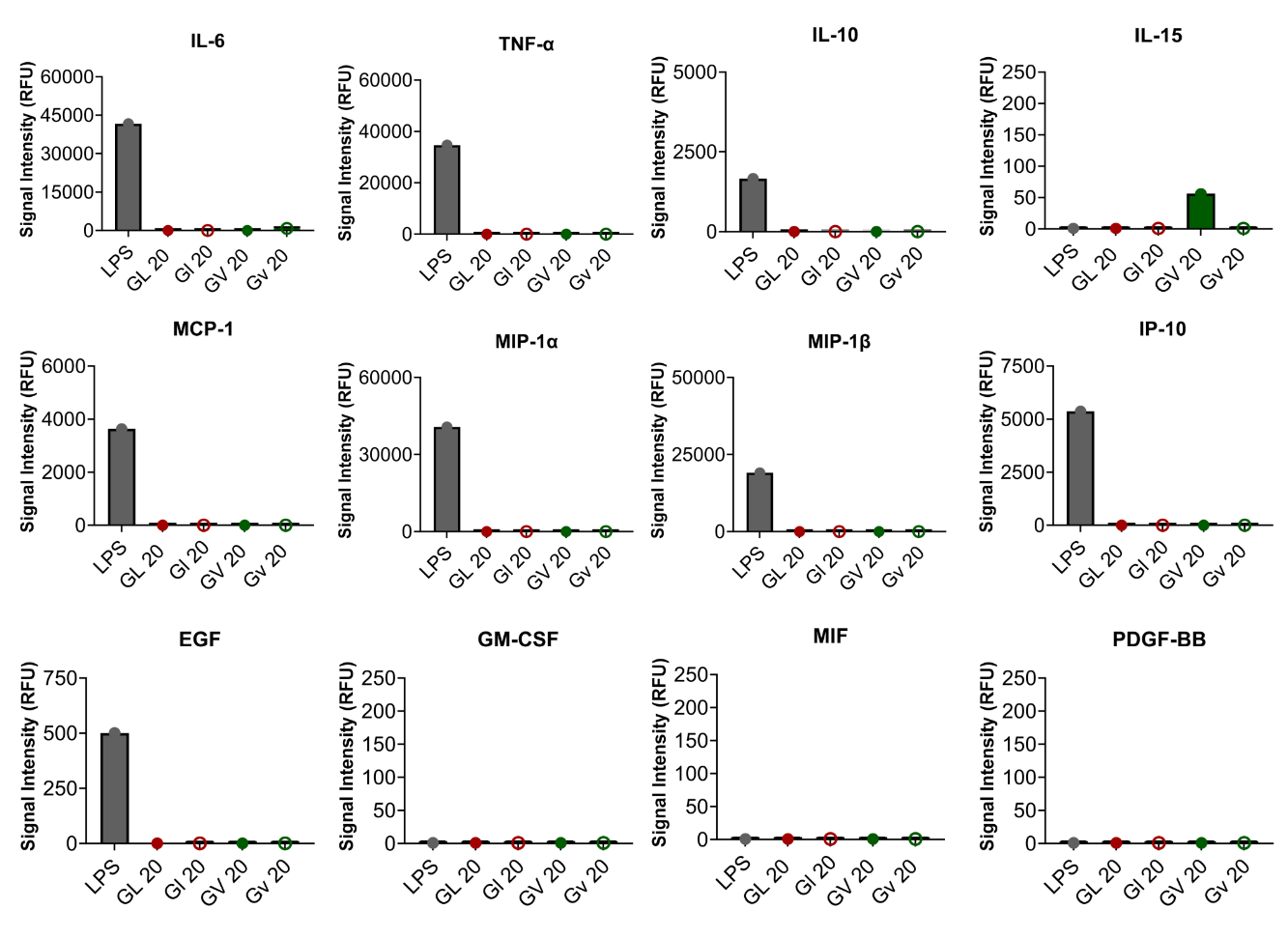

**Figure S25.** Cytokine (top row), chemokine (middle row) and (bottom row) growth factor production in BMDCs treated with L-form or D-form (GHGXY)_4_ coacervates (10 μM) for 24 h .

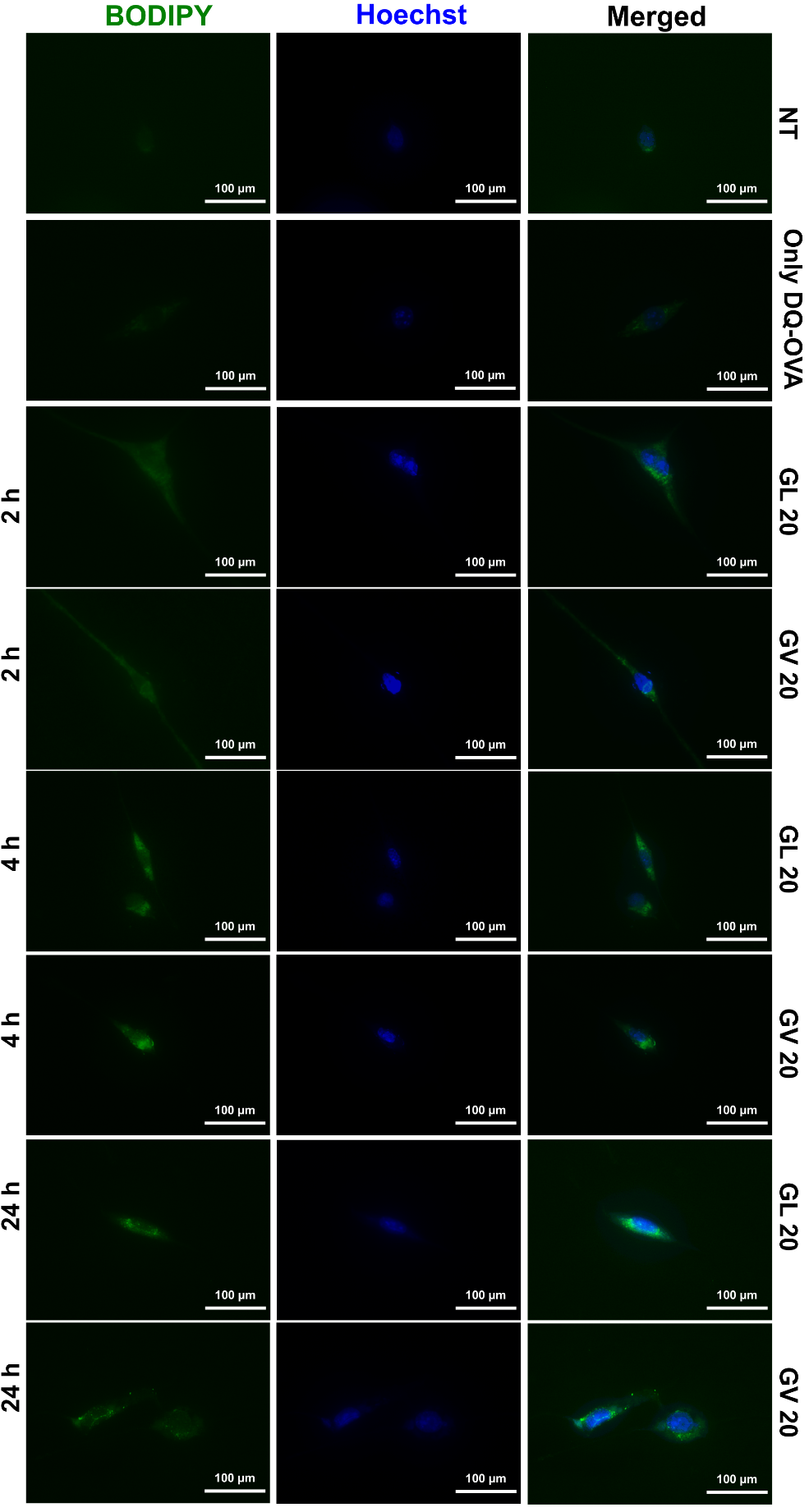
**Figure S26.** Fluoresence signal (green) from DQ-OVA processing in lysosomes following delivery with leucine or valine coacervates at different time points (2, 4, 24 h). Hoechst (blue) was used to stain the nucleus of BMDCs. Merged channels are also shown.

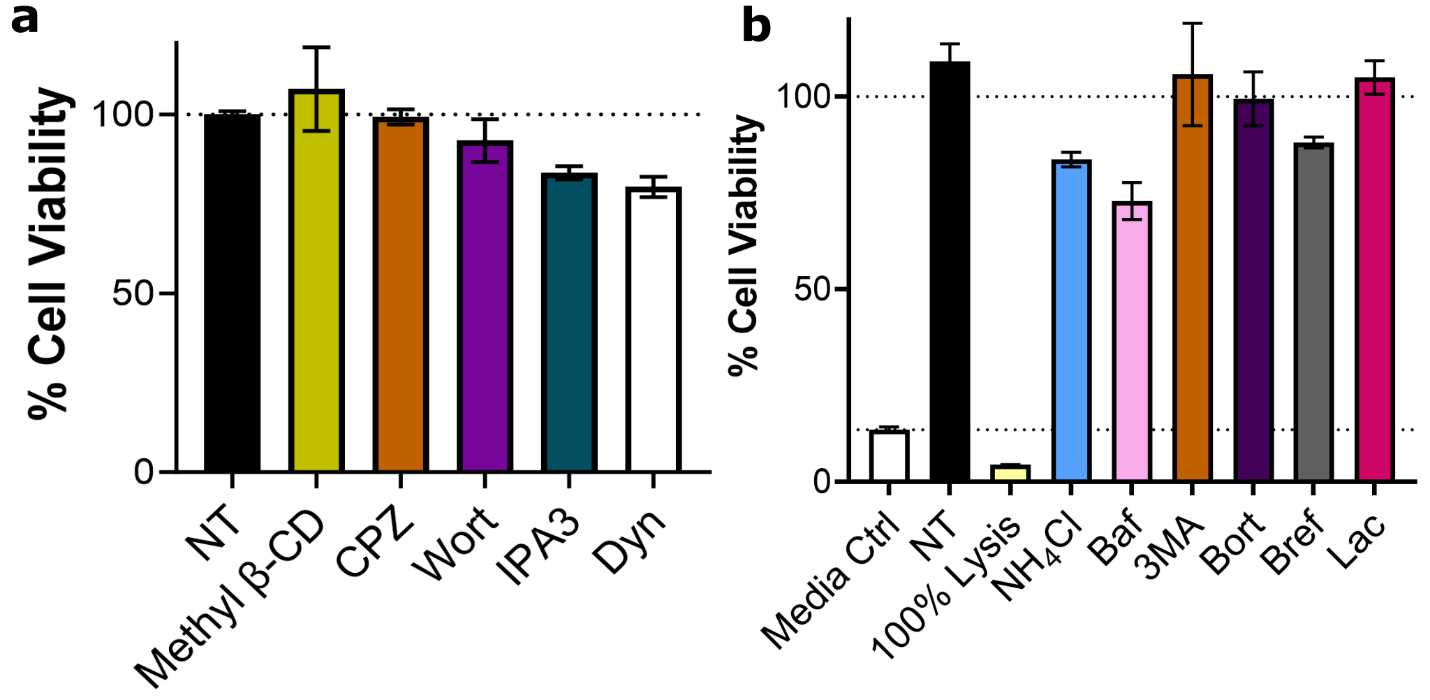

**Figure S27.** Viability of primary murine BMDCs following treatment with (a) endocytic pathway inhibitors (b) MHC class I and MHC class II pathway inhibitors at the concentrations used in the study.

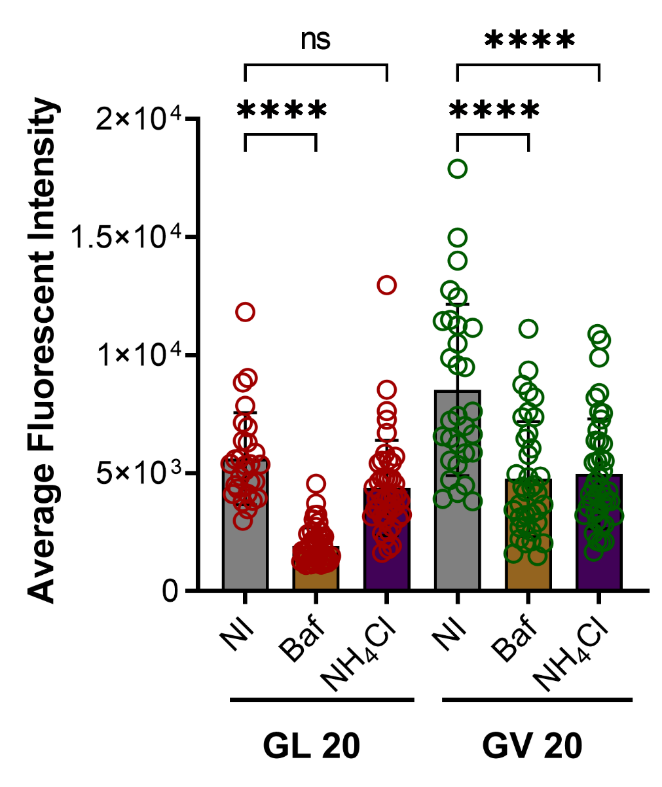

**Figure S28.** Reduction in fluorescence intensity of DQ-OVA delivered using leucine or valine coacervates in presence of Bafilomycin (Baf) or ammonium chloride (NH_4_Cl). *****p* < 0.0001 as determined by one-way ANOVA.

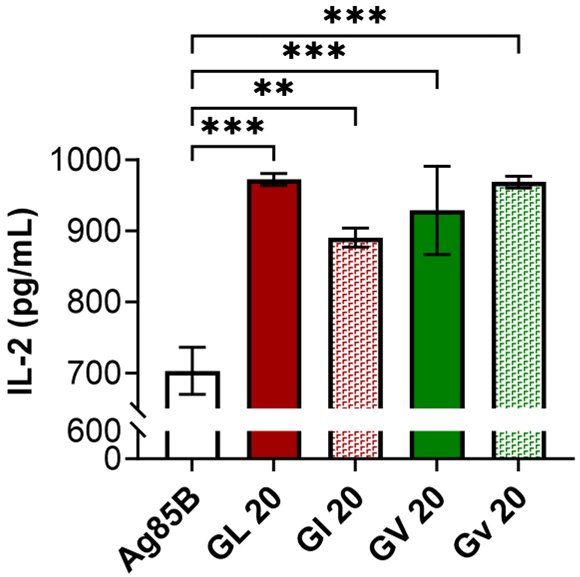

**Figure S29.** IL-2 production by BB7 hybridoma recognizing the processed Ag85B_240-254_ epitope presented in the context of MHC II following delivery using leucine or valine coacervates. *****p* < 0.0001 as determined by one-way ANOVA.

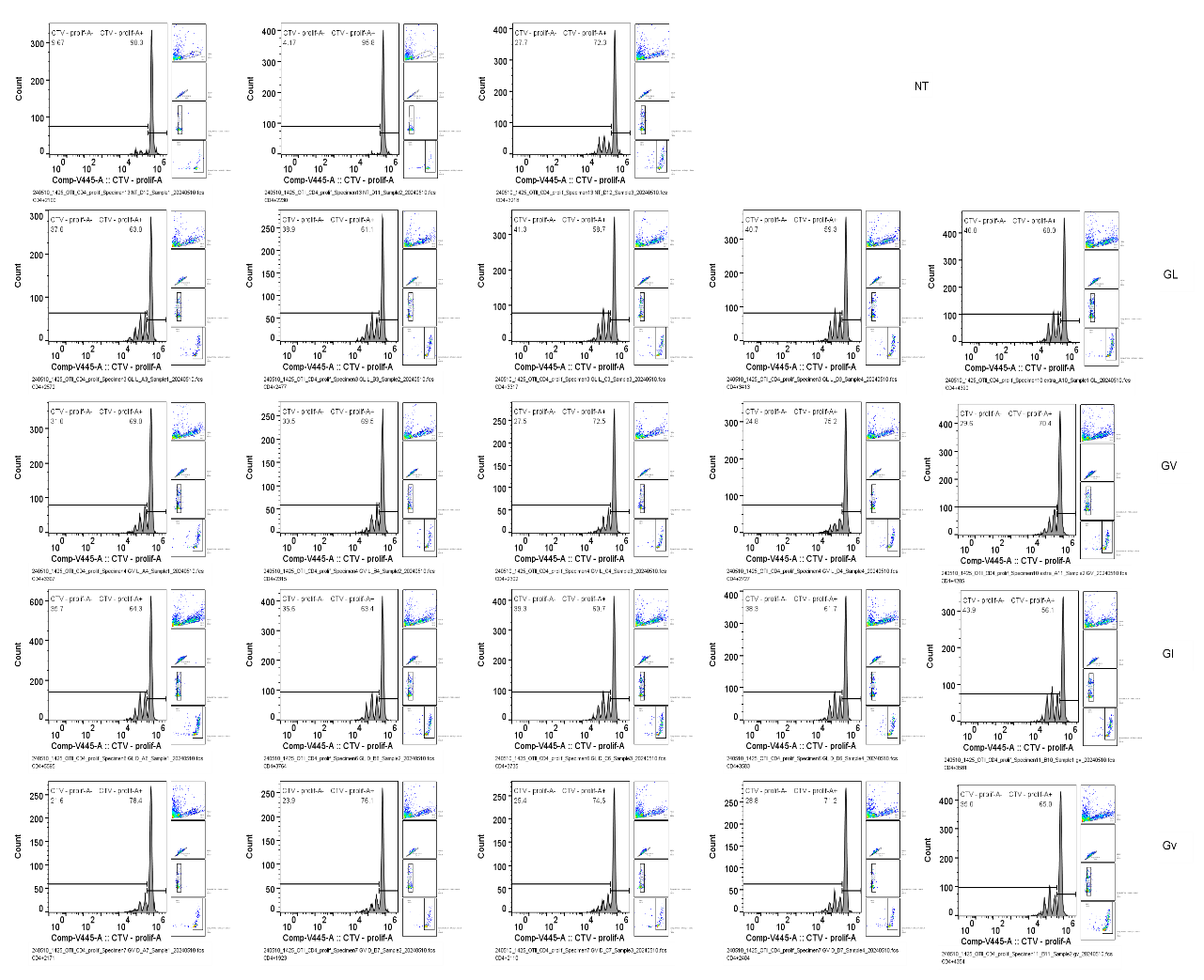

**Figure S30.** Individual histograms of CD4^+^T-cell proliferation. CD4^+^T-cells isolated from OT-II mice and labeled with CTV were overlaid on BMDCs treated with chiral (GHGLY)_4_ or (GHGVY)_4_ coacervates and proliferation measured as loss of CTV signal.

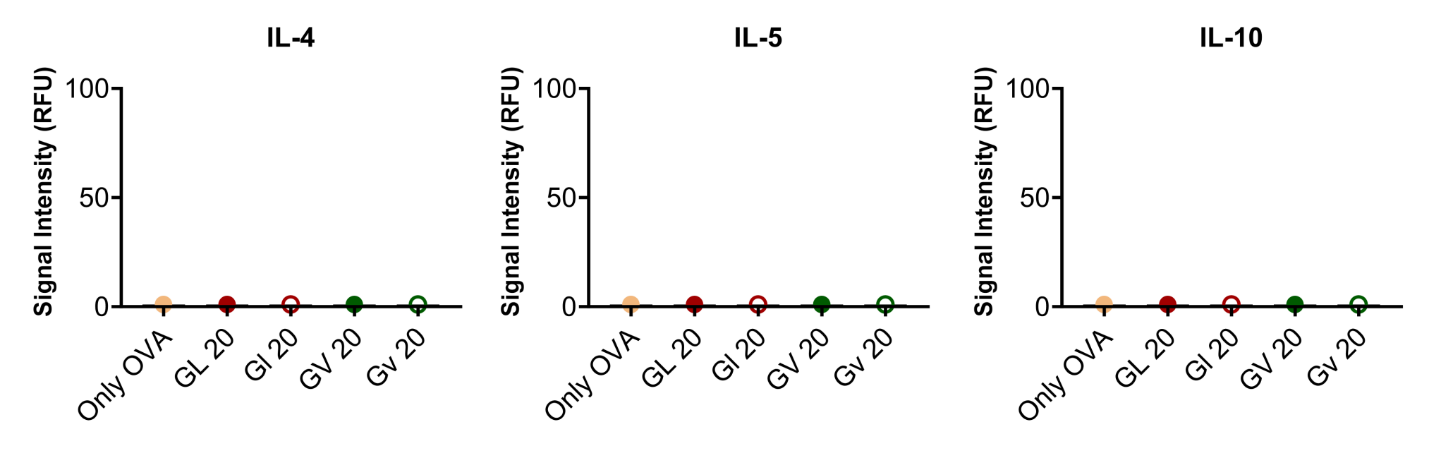

**Figure S31.** Adaptive immune responses depicting different levels of cytokines released upon treating OVA loaded L- and D- (GHGXY)_4_ droplets with antigen presenting cells (mouse BMDCs) for 24 h and overlaying with CTV-labelled OT-II mice cells for 66 h.

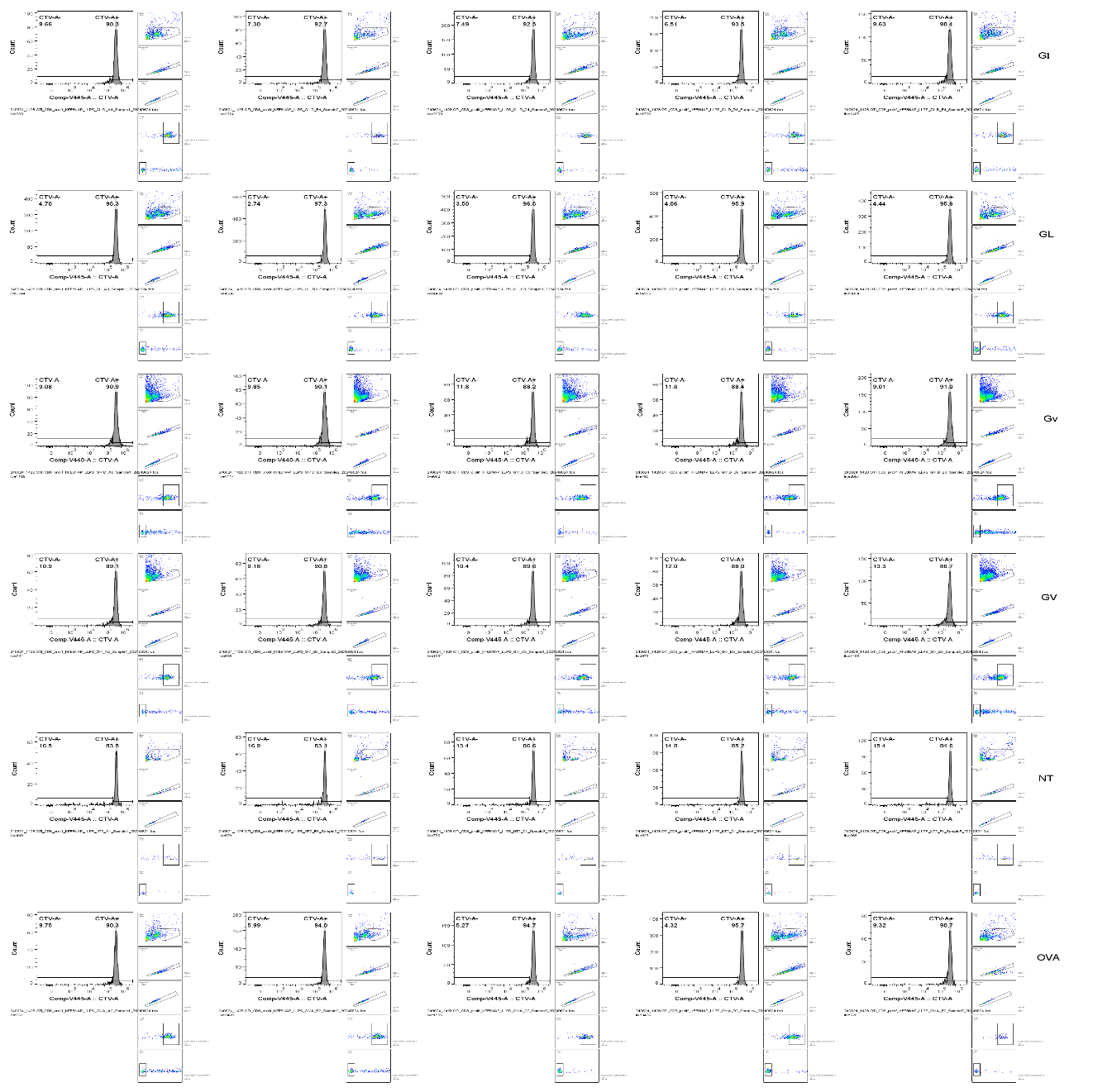

**Figure S32.** Individual histograms of CD8^+^T-cell proliferation. CD8^+^T-cells isolated from OT-I mice and labeled with CTV were overlaid on BMDCs treated with chiral (GHGLY)_4_ or (GHGVY)_4_ coacervates and proliferation measured as loss of CTV signal.

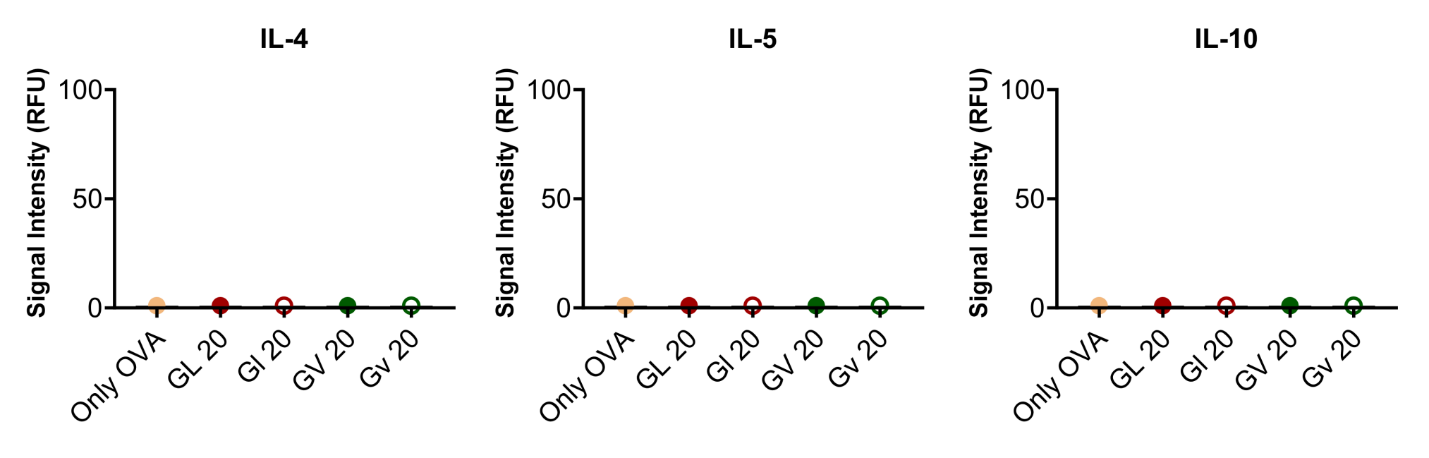

**Figure S33.** Adaptive immune responses depicting different levels of cytokines released upon treating OVA loaded L- and D- (GHGXY)_4_ droplets with antigen presenting cells (mouse BMDCs) for 24 h and overlaying with CTV-labelled OT-I mice cells for 66 h.
